# Drought and rewetting events enhance nitrate leaching and seepage-mediated translocation of microbes from beech forest soils

**DOI:** 10.1101/2020.08.03.234047

**Authors:** Markus Krüger, Karin Potthast, Beate Michalzik, Alexander Tischer, Kirsten Küsel, Florian F. K. Deckner, Martina Herrmann

**Author notes:** Corresponding author: Martina Herrmann.

## Abstract

Nitrification in forest soils is often associated with increased leaching of nitrate to deeper soil layers with potential impacts on groundwater resources, further enhanced under scenarios of anthropogenic atmospheric nitrogen deposition and predicted weather extremes. We aimed to disentangle the relationships between soil nitrification potential, seepage-mediated nitrate leaching and the vertical translocation of nitrifiers in soils of a temperate mixed beech forest in central Germany before, during and after the severe summer drought 2018. Leaching of nitrate assessed below the litter layer and in 4, 16 and 30 cm soil depth showed high temporal and vertical variation with maxima at 16 and 30 cm during and after the drought period. Maximum of soil potential nitrification activity of 4.4 mg N kg^-1^ d^-1^ only partially coincided with maximum nitrate leaching of 10.5 kg N ha^-2^. Both ammonia oxidizing bacteria (AOB) and ammonia oxidizing archaea (AOA) were subject to translocation by seepage, and AOB decreased at least by half and AOA increased by one to three orders of magnitude in their abundance in seepage with increasing soil depth. On the level of the total bacterial population, an increasing trend with depth was also observed for *Cand*. Patescibacteria while Bacteroidetes were strongly mobilized from the litter layer but poorly transported further down. Despite stable population densities in soil over time, abundances of AOA, AOB and total bacteria in seepage increased by one order of magnitude after the onset of autumn rewetting. Predicted future higher frequency of drought periods in temperate regions may result in more frequent seepage-mediated seasonal flushes of nitrate and bacteria from forest soils. Moreover, the observed translocation patterns point to taxon-specific differences in the susceptibility to mobilization, suggesting that only selected topsoil derived microbial groups are likely to affect subsoil or groundwater microbial communities and their functional potential.

## Introduction

Increasing atmospheric deposition of nitrogen (N) from anthropogenic sources has led to strongly enhanced N loads to temperate forest soils over the last decades (Borken and Matzner, 2004; Reay et al., 2008; Zhu et al., 2015). These inputs affect N cycling and support soil nitrification by elevated ammonium (NH_4_^+^) availability, increasing the risk of enhanced leaching of N compounds when the demands of N for trees and soil microbes are exceeded (Corre *et al*., 2003; Neff *et al*., 2003; Herrmann *et al*., 2005; Gundersen *et al*., 2006). In particular, NO_3_ ^-^ can rapidly be leached from soils and accumulate in subsurface compartments such as groundwater, negatively impacting its usage as drinking water supply (Gruber and Galloway, 2008; Schlesinger, 2009).

In soils NO_3_^-^ is mainly produced through nitrification driven by nitrifying prokaryotes (Leininger *et al*., 2006; Prosser, 2014; Lehtovirta-Morley, 2018). The first and rate limiting step of nitrification is the oxidation of NH_4_ ^+^, conducted by ammonia oxidizing bacteria (AOB) and archaea (AOA), followed by the oxidation of nitrite (NO_2_ ^-^) to NO_3_ ^-^ by nitrite-oxidizing bacteria (Kowalchuk and Stephen, 2001; Leininger et al., 2006). The discovery of complete ammonia oxidizers (comammox) of the genus *Nitrospira* revealed new players in this process, however, their relative contribution to soil nitrification is still under debate (Hu and He, 2017; Van Kessel et al., 2015).

Current climate models assume more frequently occurring severe summer droughts in temperate regions with critical impacts on major element cycles also in European forests (Schlesinger et al., 2016). Drought affects carbon and nitrogen cycling mediated by soil microbial processes, and decreases plant nutrient uptake (Gordon *et al*., 2008; Schlesinger *et al*., 2016; Schaeffer *et al*., 2017). Drought also reduces soil nitrification (Stark and Firestone, 1995) and N leaching by increased microbial immobilization (Schimel *et al*., 2007); it changes the soil water potential, disturbs water transport mechanics and prevents microbial mobility (Schimel, 2018). In turn, rewetting after drought can cause reactivation and rapid growth of microbes fueling nutrient cycling (Göransson et al., 2013; Placella and Firestone, 2013).

However, specific studies addressing the linkage of forest soil nitrification and NO_3_ ^-^ leaching under drought followed by rewetting conditions are still scarce.

Selective vertical translocation of functional groups of soil microbes will affect microbial community structures of the subsoil and transform its metabolic repertoire. To date, it is unknown to which extent specific functional or taxonomic groups are translocated in soils by seepage, whether nitrifying microorganisms follow depth-dependent translocation patterns, and whether their transfer is linked to changes in seepage chemistry.

Previous studies pointed to taxon-specific differential mobilization patterns of microorganism by seepage, probably linked to specific factors such as cell morphology or surface properties (Dibbern et al., 2014; Zhang et al., 2018). Transport of microorganisms in soil was mostly studied in artificial soil columns inoculated with defined microbial communities (Abu-Ashour et al., 1994; Lehmann et al., 2018), while fewer studies addressed microbial transport under field conditions (Dibbern et al. 2014, Zhang et al. 2018, Herrmann et al. 2019). But these studies did not spatially resolve the vertical translocation of microbes from the forest litter layer through the mineral horizons and how these translocation patterns vary over time, for example as a consequence of droughts or heavy rainfalls. Based on previous observations of increased microbial activity in response to rewetting after drought, we expected that rainfall periods following summer drought will result in enhanced nitrification potential, enhanced NO_3_ ^-^ leaching, and increased seepage-mediated transport of microorganisms including nitrifiers from soils of a mixed beech forest.

Thus, we aimed to (i) assess soil nitrification activity at four time points during a one-year period, which included a severe summer drought, (ii) follow NO_3_ ^-^ leaching from the litter layer to 30 cm soil depth in four depth intervals at four time points during this period, (iii) identify potential key players of soil nitrification and their susceptibility to seepage-mediated vertical translocation, and (iv) expand the analysis of translocation patterns to the total bacterial community.

## Materials and methods

### Sampling site and sampling procedure

Soil and seepage samples were obtained from a mixed beech forest at the Hainich Critical Zone Exploratory (CZE) in western Thuringia, Germany, which was established within the framework of the Collaborative Research Center (CRC) AquaDiva (Küsel et al., 2016). The CZE spans a 6 km transect along a hill slope with managed deciduous forest as the primary land use type at the hilltop. This forest is dominated by European beech (*Fagus sylvatica*) with minor contribution of sycamore maple (*Acer pseudoplatanus*) and common ash (*Fraxinus excelsior*) (Potthast et al., 2017). Calcaric Cambisol is the dominating soil type of this mixed beech forest (Potthast et al., 2017). Zero-tension lysimeters (300 cm_2_ sampling area) were installed within defined plots of the forested area (Potthast et al., 2017). These lysimeters allow sampling of soil leachate (seepage) from defined depth intervals by collecting seepage that percolated through the litter layer, and through litter layer plus the upper 4 cm, 16 cm, and 30 cm of the soil, respectively (Figure 1). In addition, on site soil moisture and soil temperature data were recorded in ten-minute intervals by moisture sensors at 4 cm, 16 cm and 30 cm depth and temperature sensors at 4 cm and 16 cm depth at one study plot. Sampling of seepage was conducted four times in 2018 (February, May, July, November) and additionally in March 2019. In summer 2018 a severe drought reduced precipitation to 55% from April to September (half year of 2018: 191 mm) compared to former years (half year average of 2015 - 2019: 343 mm) recorded by the Heuberg weather station of the Hainich CZE (Lehmann and Totsche, 2020). Seepage was sampled at two lysimeter plots with two replicates of each lysimeter depth (litter layer, 0 to 4 cm, 0 to 16 cm, and 0 to 30 cm). In addition, soil samples were taken in three replicates at 5 cm and 10 cm depth nearby the corresponding lysimeters. Soil samples for later DNA extraction were frozen immediately on dry ice and stored at −80°C. Soil samples for the determination of potential nitrification rates and physicochemical analysis as well as seepage samples were stored at 4°C until further processing within 24 hours. For molecular analysis, seepage samples were filtered onto 0.2 µm pore size polyethersulfone filters (Supor, Pall Corporation, USA) and kept frozen at −80°C until DNA extraction. Filtrates were used for later analysis of inorganic nitrogen.

**Figure 1.**
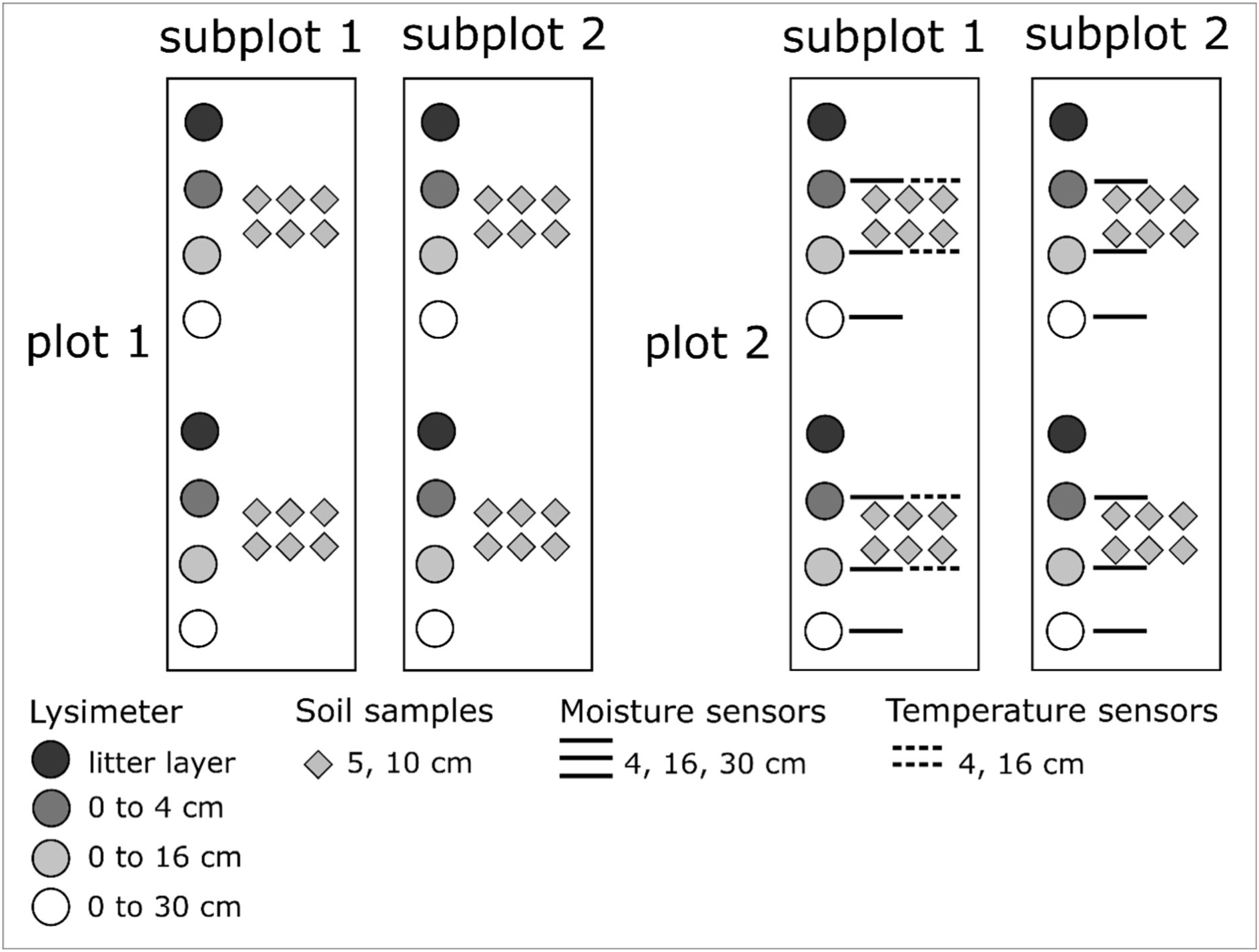
Plot design of lysimeters installed below the litter layer, 0 - 4 cm, 0 −16 cm and 0 - 30 cm depth intervals at a mixed beech forest in the Hainich region (Germany). Two similar designed plots consisting of two subplots with two replicates from each lysimeter depth. Soil samples from 5 cm and 10 cm were taken nearby corresponding lysimeters. Additional sensors for measuring soil moisture at 4, 16 and 30 cm, and temperature at 4 and 16 cm were available.

### Potential soil nitrification activity

The potential nitrification activity of soils was determined by slurry incubations (Hart et al., 1994; Yao et al., 2011). 15 g of fresh soil from 5 and 10 cm depth was transferred to 250 ml Erlenmeyer flasks and mixed with 100 ml sterile NH_4_Cl solution (0.5 mM). Slurry samples were incubated aerobically at 15°C in the dark with constant shaking (150 rpm) for 5 days. Subsamples of 1.8 ml were taken every 24 hours, centrifuged briefly for sedimentation of soil particles, and the supernatant was stored at −20°C for later measurement of NO_3_ ^-^ concentrations. Potential nitrification rates were calculated from the slope of the increase of NO_3_ ^-^ concentrations over time.

### Physicochemical properties of soil and seepage

Soil water content was determined by drying the samples over 24 hours at 105°C. Further analysis of soil chemical parameters was conducted with soil suspended in deionized water at a ratio of 1:2.5. pH of both soil and seepage samples was measured with a glass electrode. In seepage samples total organic carbon (TOC), total nitrogen (TN) as well as dissolved organic carbon (DOC) and dissolved nitrogen (DN) concentrations (<0.45 µm) were determined by catalytic thermal oxidation and NDIR/ chemoluminescence detection, respectively (Shimadzu, Germany). Particulate organic carbon (POC) and particulate nitrogen (PN) were calculated from the difference between total and dissolved concentrations. Concentrations of NH_4_ ^+^ and NO_3_ ^-^ were determined photometrically using standard colorimetric approaches (Scheiner, 1974; Seelmeyer, 1954). Fluxes of NO_3_ ^-^ and NH_4_ ^+^ were calculated from the concentration of the respective compound in the seepage samples multiplied by the seepage volume and the lysimeter area and expressed in mg N m^-2^

### Extraction of DNA and quantitative PCR

Genomic DNA was extracted using the DNEasy PowerSoil kit (Qiagen, Germany) according to the manufacturer’s instructions from 0.25 g soil or seepage filter material and stored at - 20°C. Abundances of bacterial and archaeal *amoA* genes and bacterial 16S rRNA genes were determined by quantitative PCR using a Mx3000P instrument (Agilent, CA). Primer combinations were AmoA-1F/ AmoA2R (Rotthauwe *et al*., 1997) for bacterial *amoA*, Arch-AmoAF/ Arch-AmoAR (Francis et al., 2005) for archaeal *amoA* and Bac8fmod/Bac338Rabc for bacterial 16S rRNA gene (Daims et al., 1999; Loy et al., 2002), with cycling conditions reported previously (Herrmann *et al*., 2012). Reactions were set up in a 25 µl total volume with 12.5 µl Brilliant SYBR Green qPCR Master Mix (Agilent), 0.4 µM of each primer and 5 µl of prediluted DNA. Standard curves were generated by serial dilutions of standards containing mixtures of plasmid DNA containing the respective gene as cloned insert. Standard curves were linear from 5 x 10^8^ to 50 copies per reaction for *amoA* genes and from 5 x 10^8^ to 5 x 10^2^ copies per reaction for bacterial 16S rRNA genes, respectively, with R_2_ > 0.99 and qPCR efficiencies ranging from of 80 - 95%.

### Illumina MiSeq amplicon sequencing

Libraries for amplicon sequencing of bacterial 16S rRNA genes were prepared using a two-step barcoding approach. The primer combination Bakt341F/ Bakt785R covering the V3 to V4 region of the bacterial 16S rRNA gene (Herlemann et al., 2011) was used in a first PCR. The primers were modified with an adaptor overhang for Illumina sequencing, allowing barcoding through a second PCR step. In the first step genomic DNA of soil and seepage samples was amplified in triplicates with 1 - 5 ng DNA in a 15 µl reaction volume, 0.4 µM of each primer, 1 µg/ µl BSA and 10X HotStartTaq Master Mix (Qiagen). Cycling conditions included initial denaturation at 95°C for 15 min, 30 cycles at 94°C for 45 sec, 55°C for 45 sec, 72°C for 45 sec and final elongation at 72°C for 10 min. The size and integrity of PCR products was confirmed by agarose gel electrophoresis. Sequencing was performed on an Illumina MiSeq platform using v3 chemistry.

### Sequence analysis

Analysis of bacterial 16S rRNA amplicon reads were performed using Mothur v1.39 (Schloss et al., 2009), following the MiSeq-SOP (Kozich et al., 2013). Quality trimming of reads was done by excluding those reads with more than 8 homopolymers and sequences which include ambiguous bases. Remaining sequences were pre-clustered allowing for 1 base difference per 100 bp, followed by chimera search using the uchime algorithm implemented in Mothur (Edgar et al., 2011). Sequences were taxonomically classified using the SILVA reference data base release 132 (Quast et al., 2013). Non-bacterial lineages were removed, and remaining sequences were clustered with a 0.03 cutoff for OTU assignment. Further downstream sequence analysis was performed using the R package phyloseq (McMurdie and Holmes, 2013). Sequence data generated in this study was submitted to the European Nucleotide Sequence Archive (ENA) with the project number PRJEB39355, accession numbers ERS4817796 – ERS4817857.

### Estimation of fluxes of microbial cells and their contribution to carbon and nitrogen fluxes

We estimated which fraction of TOC and POC as well as of TN and PN in seepage was accounted for by bacterial cells. We assumed a mean carbon content of 39 fg per cell and a mean nitrogen content of 12 fg per cell under C-limited conditions (Vrede et al., 2002) for bacterial cells. For *Cand*. Patescibacteria we considered a lower mean carbon content of 0.7565 fg per cell and a mean nitrogen content of 0.233 fg per cell due to a 51 times smaller cell volume of these ultra-small cells (Luef et al., 2015) compared to cell volumes given in Vrede et al. (2002). The number of exported cells was estimated on family-level by first multiplying 16S rRNA gene abundances per liter by the seepage volume collected at a given time point and by the relative abundance of a given family obtained from Illumina amplicon sequencing. These numbers were then corrected by the family specific average 16S rRNA operon number obtained from the ribosomal RNA operon database (rrnDB: https://rrndb.umms.med.umich.edu/) (Stoddard et al., 2015), to obtain accurate estimates of cell abundances. Results of these family-based calculations were summed up to estimate the total number of bacterial cells translocated by seepage per lysimeter and event. These numbers of estimated bacterial cells were then multiplied with the assumed cell carbon and nitrogen content, and the resulting estimate of microbial cell carbon and nitrogen was compared to the content of TOC, POC, TN and PN in the respective sample.

Microbial cell fluxes were estimated based on the calculations above for the total bacterial population for samples obtained in February, July and November 2018. An average operon number of 2.7 was used to correct 16S rRNA gene abundances from March 2019 where no sequence data was available. In similar calculations, we assumed an average operon number of 2.5 for AOB-*amoA* (Norton et al., 2002), and one gene copy per cell for AOA-*amoA* (Hallam et al., 2006). Estimated cell fluxes were then calculated as described above for NO_3_ ^-^ and NH_4_ ^+^ fluxes and were expressed in cells m^-2^.

To estimate the fraction of translocated nitrifiers by seepage from the litter layer to 30 cm depth relative to their abundance in the soil volume through which seepage had percolated, we first estimated how many AOB and AOA cells were transported via seepage through 30 cm of soil using the same approach to calculate cell fluxes as described above. Numbers of ammonia oxidizer cells in the corresponding soil volume were estimated by extrapolating estimated cell abundances per g soil to cell abundances in the total soil volume from which seepage was collected, assuming a soil volume of 8.5 L with an average density of 1.03 g cm^-3^. The exported fraction of cells was obtained by dividing the estimated abundances of cells in seepage by the estimated abundance of cells in the corresponding soil volume. This approach yields conservative estimates as microbial abundances in 10 cm soil were used to extrapolate total abundances across the upper 30 cm of soil. Since microbial abundances in soil most likely decrease with increasing soil depth, the fraction of cells mobilized by seepage might be even higher.

### Statistical analysis

All statistical tests were performed in the R environment (v.3.6.2) and RStudio (v1.2.5033). Correlations between physicochemical parameters and microbial gene abundances were performed using Spearman’s rank correlation. Non-parametric statistics were used to test for differences between means of soil and seepage parameters as well as microbial gene abundances. As global test of significance, the Kruskal-Wallis rank sum test was applied, followed by Dunn’s multiple comparison test between samples. Principal coordinate analysis (PCoA) based on Bray-Curtis distance matrices was used to cluster soil and seepage samples based on their similarity/ dissimilarity. Permutational multivariant analysis of variance (PERMANOVA) of factors accounting for OTU variances across sampling months and depths was performed using Adonis functions from R package vegan (Oksanen et al., 2008). Default 999 permutations were assessed based on Bray-Curtis distance.

## Results

### Potential nitrification activity and nitrifier abundance in forest soils

Soil potential nitrification rates (PNR) of the mixed beech forest showed temporal heterogeneity between samples ranging from < 0.1 to 4.4 µg N g dw^-1^ d^-1^ (Figure 2) with maximum values in July and lowest PNR in November. Differences in PNR between 5 and 10 cm soil depth were minor and only apparent by trend for samples taken in May and November.

**Figure 2.**
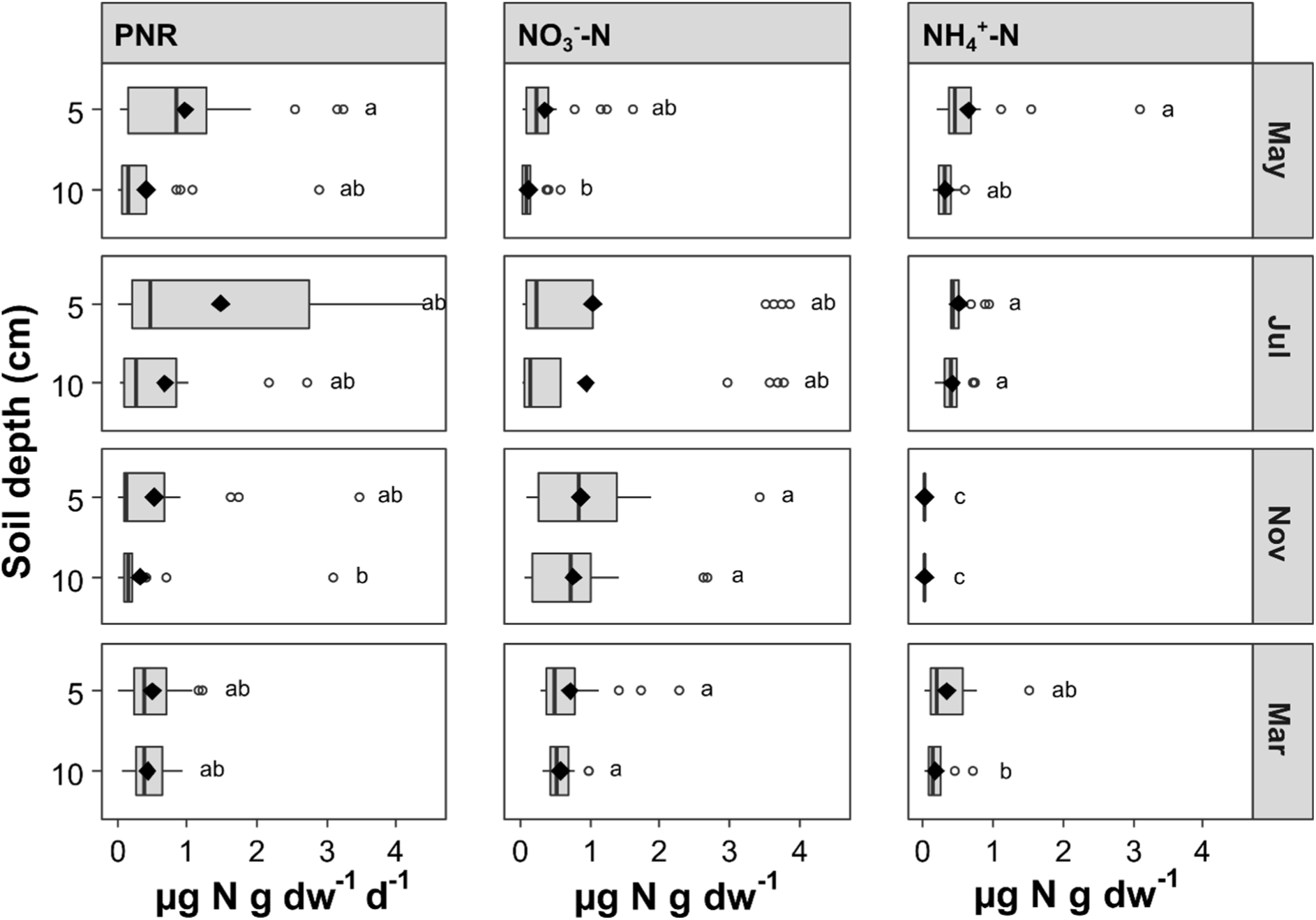
Soil potential nitrification rate (PNR), nitrate-N content and ammonium-N content in 5 cm and 10 cm depth at mixed beech forest plots. Soil samples were taken in May, July, November 2018, and March 2019. Individual boxplots summarize the measured parameters at particular depths and black diamonds represent mean of respective data. Boxplots show lowest observation, 25% quartile, median, 75% quartile, highest observation, and outlier as white dots. The lowercase letter code indicates significant differences based on Dunn’s multiple comparison test (*p* < 0.05).

The observed spatial and temporal patterns of PNR were reflected by soil concentrations of NO_3_ ^-^ with maxima of 3.9 µg N g dw^-1^ and 3.4 µg N g dw^-1^ in July and November, respectively (Figure 2). NO_3_ ^-^ content of soils in May and March was lower with maxima of 1.6 µg N g dw^-1^ and 2.3 µg N g dw^-1^, respectively. Similarly, NH_4_ ^+^ content of soils displayed differences between sampling months with a maximum of 3.1 µg N g dw^-1^ in May and low values of less than 0.1 µg N g dw^-1^ in November (Figure 2). Significant differences in both NO_3_ ^-^ and NH_4_ ^+^ content between 5 and 10 cm soil depth were observed in May and March.

In addition, we assessed environmental conditions of nitrogen formation by the determination of soil moisture and temperature for every sampling time point (Table S1). Across all sites and time points, PNR were positively correlated with soil NO_3_ ^-^ content (R = 0.28, *p* < 0.001) and NH_4_ ^+^ content, but not directly with soil temperature and moisture (Table S2).NH_4_ ^+^ content showed a positive correlation to soil water content (R = 0.36, *p* < 0.01), while the correlation between NO_3_ ^-^ and soil water content was negative (R = −0.31, *p* < 0.01).

During spring and summer 2018 an extensive summer drought led to long periods of low precipitation and low soil moisture. Among the sampling time points water content was lower in July and November with an average of 23% water content compared to May 2018 and March 2019 and samples with an average of 40%, while soil temperature in July was highest with around 9°C. Consequently, maximum PNR in July coincided with highest soil temperature and lowest soil moisture.

Soil PNR was also positively correlated with abundances of bacterial *amoA* genes as a measure of the abundance of ammonia-oxidizing bacteria in the soil (R = 0.56, *p* < 0.0001) (Table S2). Moreover, AOB-*amoA* gene abundance showed a significant positive correlation with soil NO_3_ ^-^ content (R = 0.34, *p* < 0.01) and pH (R = 0.62, *p* < 0.001). To test whether the observed PNR could sufficiently be explained by activities of the detected AOB populations, we estimated the per cell activity of AOB by dividing the PNR by the bacterial *amoA* abundance for a given sample, assuming an average of 2.5 *amoA* gene copies per cell (Norton et al., 2002). Resulting mean estimated per cell activities were 11.7 fmol N cell^-1^ h^-1^ in May, 24.5 fmol N cell^-1^ h^-1^ in July, 5.4 fmol N cell^-1^ h^-1^ in November and 2.6 fmol N cell^-1^ h^-1^ in March, which, except for the values in May, are similar to previously reported cell activities of 2 to 7 fmol N cell^-1^ h^-1^ (Lu et al., 2015; Taylor and Bottomley, 2006). Estimated mean cell activities of AOA ranged from 4.9 fmol N cell^-1^ h^-1^ in July up to 31.4 fmol N cell^-1^ h^-1^ in November exceeding documented values at least ten times (Könneke et al., 2005; Lu et al., 2015).

Across all sites and time points, bacterial *amoA* and archaeal *amoA* genes ranged from 2.5 x 10^3^ to 1.4 x 10^8^ g dw^-1^ and 6.7 x 10^3^ to 4.7 x 10^7^ g dw^-1^, respectively, without significant differences between months or soil depths (Figure 3). In contrast, bacterial 16S rRNA gene abundances showed minor significant changes between 5 cm and 10 cm depth between the months ranging from 4.3 x 10^8^ to 6.6 x 10^10^ g dw^-1^.

**Figure 3.**
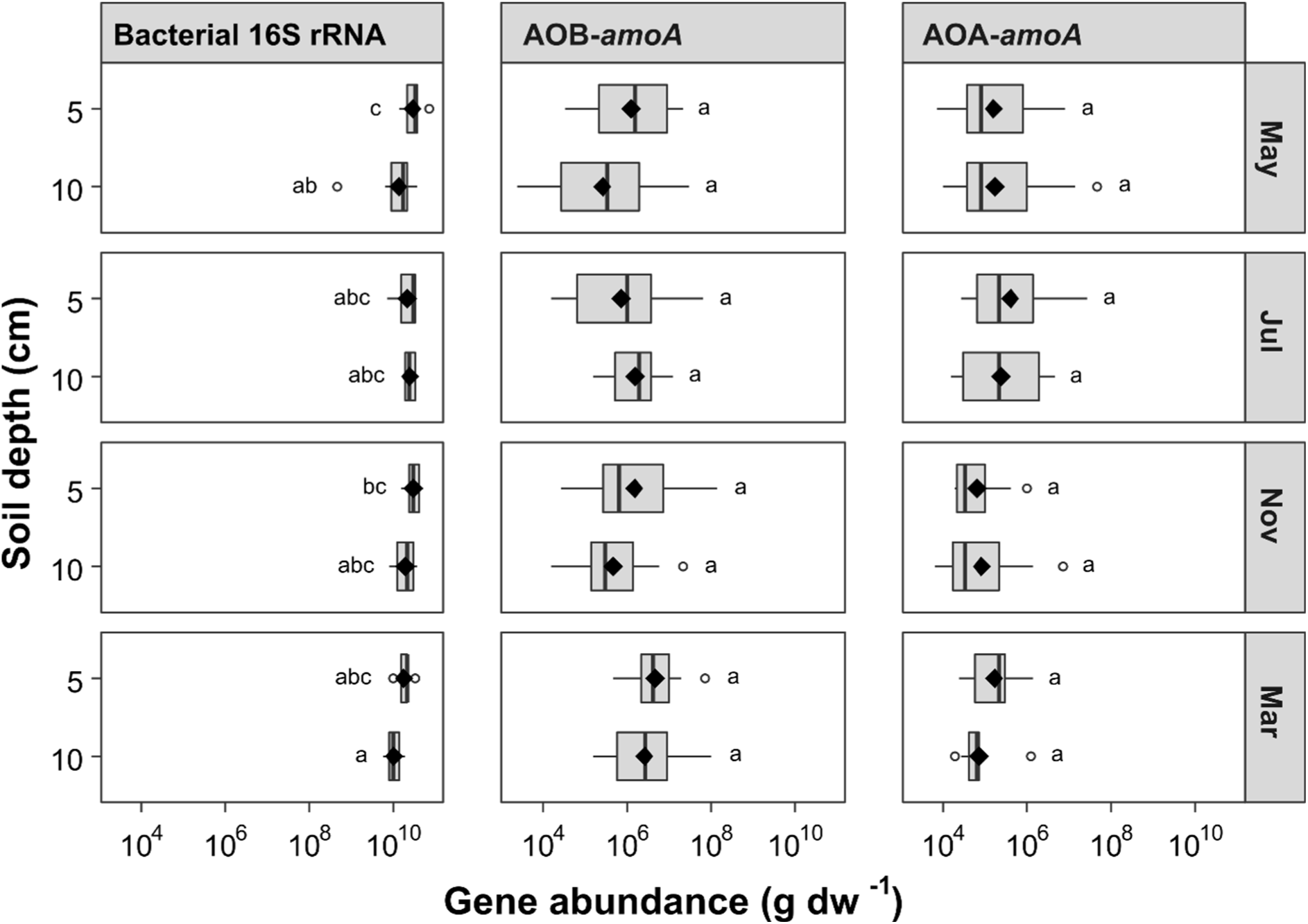
Microbial gene abundance of 16S rRNA, bacterial *amoA* and archaeal *amoA* in soil depths of 5 cm and 10 cm at mixed beech forest plots displayed in genes per gram dry weight. Soil samples were taken in May, July, November 2018, and March 2019. Individual boxplots summarize the gene abundances at particular depths and black diamonds represent mean of respective data. The lowercase letter code indicates significant differences based on Dunn’s multiple comparison test (*p* < 0.05).

### Leaching of nitrate from forest soils

The vertical export of NO_3_ ^-^ varied strongly across sampling depths and months. Highest NO_3_ ^-^ fluxes were observed in July and November with maxima of 875 mg N m^-2^ and 1051 mg N m^-2^, respectively, while seepage in February and March showed lower NO_3_ ^-^ fluxes with maxima of 342 mg N m^-2^ and 423 mg N m^-2^, respectively (Figure 4). In July and November, NO_3_^-^ fluxes increased from the litter layer towards 4 cm and 16 cm depth and then decreased again, whereas in March 2019, NO_3_^-^ fluxes increased continuously from 125 mg N m^-2^ below the litter layer to 350 mg N m^-2^ at 30 cm soil depth. In February, NO_3_^-^ fluxes showed only low vertical variation.

**Figure 4.**
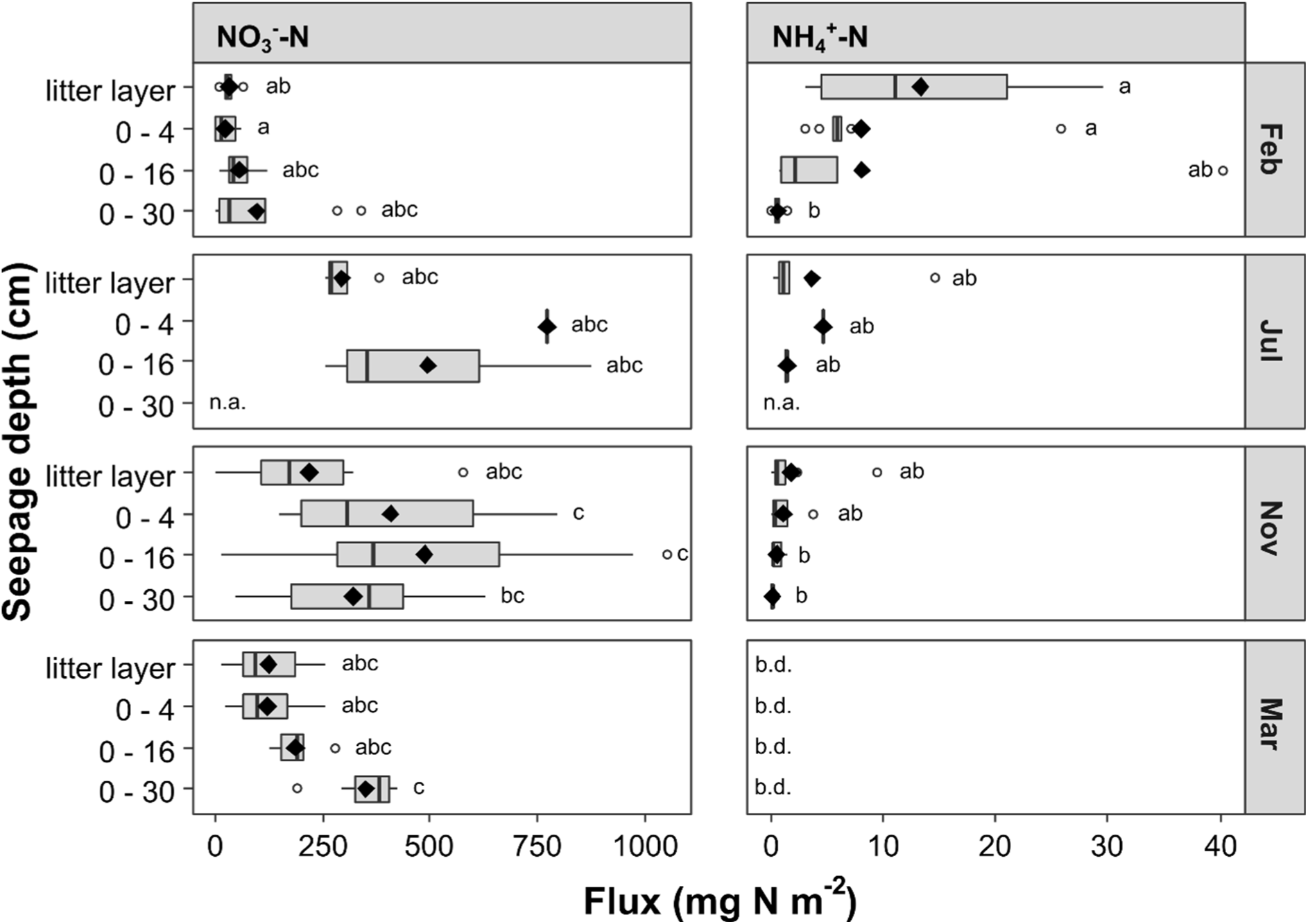
Calculated fluxes in seepage of N-nitrate and N-ammonium across four lysimeter depths of below the litter layer, 0 - 4 cm, 0 - 16 cm and 0 - 30 cm soil depth at mixed beech forest plots. Seepage samples were taken in February, July, November 2018, and March 2019. No seepage was available from 0 to 30 cm interval in July caused by low precipitation during summer drought. Individual boxplots summarize nitrogen fluxes at particular depths and black diamonds represent mean of respective data (n.a. = no sample available, b.d. = below detection limit of 1.4 µg N L^-1^). The lowercase letter code indicates significant differences based on Dunn’s multiple comparison test (*p* < 0.05).

Seepage-mediated fluxes of NH_4_ ^+^ were 10 to 100 times lower than NO_3_^-^ fluxes across all depths and months (Figure 4) and showed contrasting vertical trends with a negative correlation to NO_3_ ^-^ fluxes across all samples (R = −0.25, *p* < 0.05). Maximum NH_4_ ^+^ fluxes were observed in February with 17 mg N m^-2^ when NO_3_ ^-^ fluxes were lowest, while other months showed less than 10 mg N m^-2^, and NH_4_ ^+^ concentrations in seepage even remained below the detection limit of 0.05 mg N m^-2^ in March 2019. Across all sites, NH_4_ ^+^ and NO_3_ ^-^ fluxes were positively correlated with soil temperature (R = 0.64, *p* < 0.05 and R = 0.65, *p* < 0.01).

### Translocation of nitrifiers by seepage

Following the analysis of patterns of inorganic nitrogen leaching, we investigated to what extent ammonia-oxidizing bacteria and archaea as well as the total bacterial population were also subjected to seepage-mediated translocation. In February 2018, estimated bacterial fluxes by seepage ranged from 7.2 x 10^8^ to 1.4 x 10^11^ cells m^-2^ and showed a slight decrease towards 30 cm depth (Figure 5). A similar vertical trend was observed in November and March with one order of magnitude higher abundances compared to February and July. Ammonia oxidizers fluxes in February ranged from 2.2 x 10^5^ to 1.3 x 10^8^ cells m^-2^ for AOB and from 5.4 x 10^4^ to 2.7 x 10^7^ cells m^-2^ for AOA (Figure 5). Similar to the total bacteria population, bacterial and archaeal ammonia-oxidizers were 10 times more abundant in seepage in November and March.

**Figure 5.**
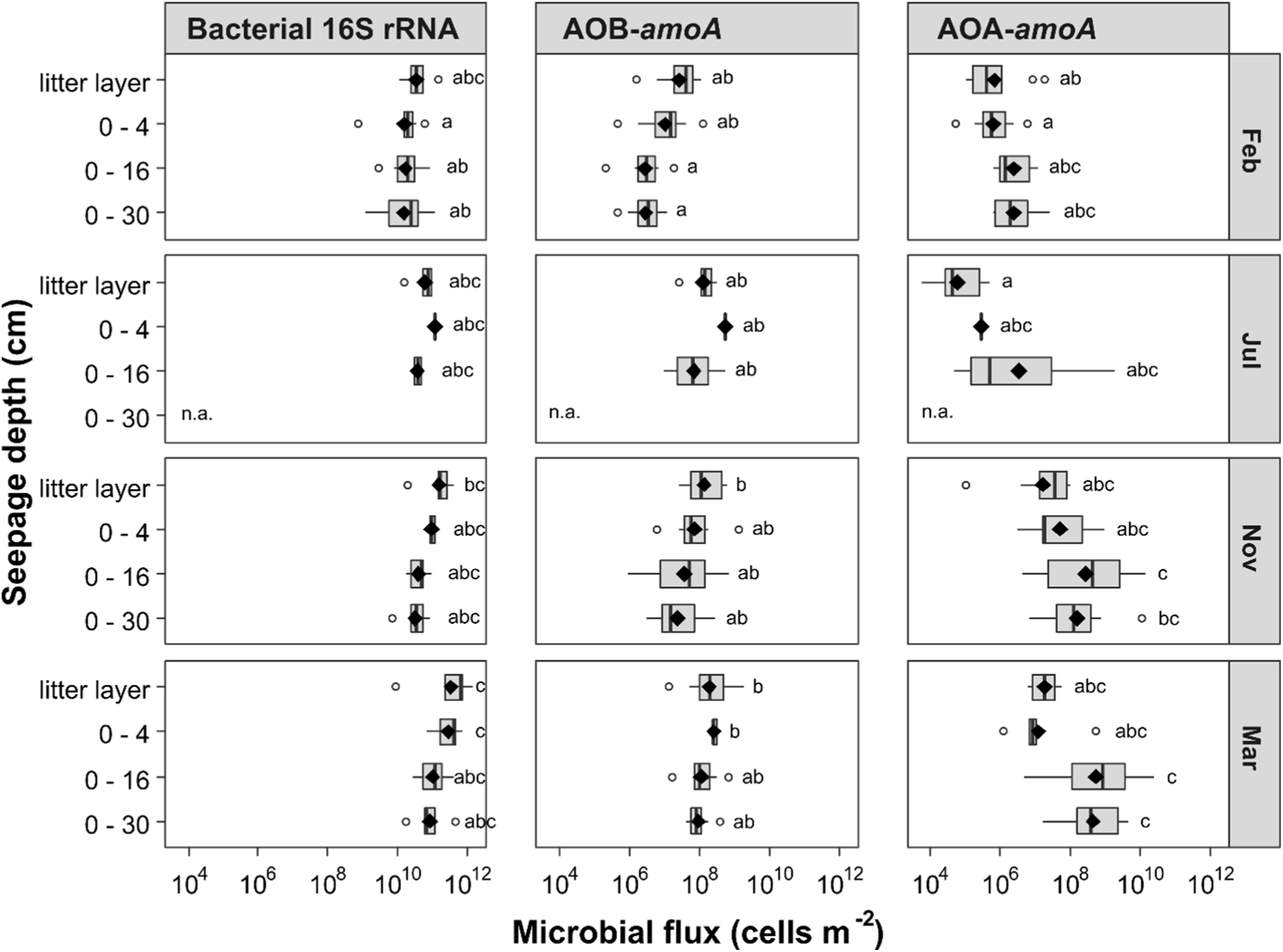
Microbial cell flux per m^-2^ of bacteria, bacterial ammonia-oxidizers (AOB) and archaeal ammonia-oxidizers (AOA) of seepage after percolation of the water through the litter layer, 0 - 4 cm, 0 −16 cm and 0 −30 cm of soil at mixed beech forest plots. Cell counts were derived from gene quantification. Seepage samples were taken in February, July, November 2018, and March 2019. Individual boxplots summarize the gene abundances at particular depths and black diamonds represent mean of respective data (n.a. = no sample available). The lowercase letter code indicates significant differences based on Dunn’s multiple comparison test (*p* < 0.05).

The two groups of ammonia oxidizers showed an opposing trend of seepage-mediated vertical translocation. While AOB fluxes were higher right below the litter layer compared to 30 cm depth, AOA fluxes were higher in seepage obtained at 16 cm and 30 cm sampling depth. Estimations of the fractions of ammonia oxidizers in the total microbial population based on *amoA*/16S rRNA gene ratios revealed that both AOB and AOA accounted for a larger fraction of the microbial community in seepage (Figure S1) than in soil (Figure S2). In addition, AOA-*amoA*/16S rRNA gene ratios in seepage increased towards 16 and 30 cm depth compared to seepage collected below the litter layer. AOA*-amoA*/AOB-*amoA* gene ratios revealed that AOB outnumbered AOA by one to two orders of magnitude in soils at 5 and 10 cm depth but this ratio was strongly altered in seepage with increasing depth, resulting in similar abundances of the two groups of ammonia oxidizers or even higher abundances of AOA in seepage collected at 30 cm depth.

Further, we estimated the fraction of ammonia oxidizers mobilized by the percolating water compared to the assumed abundance of ammonia oxidizers in the total soil volume which was percolated by seepage. For seepage collected at 30 cm depth, our estimations suggest that 1% of the AOB cells and 12% of the AOA cells presumably present in the soil were mobilized by seepage, providing further support for a stronger vertical translocation of AOA compared to AOB to soil compartments below 30 cm.

### Estimated bacterial contribution to carbon and nitrogen fluxes

Carbon and nitrogen in microbial cells contribute to the total fluxes of carbon and nitrogen. Bacterial cells accounted for an estimated average of 3% and 1.5% of the TOC and TN, respectively, in seepage (Figure S3). We tested the relationship between bacterial cell numbers and fluxes of TOC and TN across all sites and time points and found a significant positive correlation of the bacterial cell numbers with TOC (R = 0.29, *p* < 0.05) and TN (R = 0.49, *p* < 0.001) but no significant correlation of estimated cell fluxes with POC and PN. Bacterial cells made up to 30% of the POC and PN exported by seepage but these proportions differed substantially among months and depths where seepage was sampled. Approximately 20% of the POC in seepage, especially below the litter layer and in 4 cm depth, was accounted for by bacterial cells in July and November compared to only 5% in February (Figure S3). This estimated fraction decreased with depth in July and November, while the fraction was highest towards 30 cm depth in February. For nitrogen, we estimated that 21%, 34%, and 17% of PN below the litter layer were accounted for by bacteria in February, July, and November, respectively, followed by a decrease of this estimated fraction with increasing depth.

### Patterns of seepage-mediated translocation of the bacterial community

A total of 67081 OTUs were obtained after sequence analysis of the bacterial 16S rRNA gene. The nitrifying community represented by the three genera *Nitrospira, Nitrosospira* and *Nitrosomonas* made up less than 1% of the total bacterial community in seepage and soil (Figure 6), which agreed with the numbers obtained from *amoA*- and 16S rRNA gene targeted qPCR. *Nitrospira* showed the highest relative proportion among nitrifiers in seepage followed by *Nitrosospira* and *Nitrosomonas*. Following the trends observed by *amoA* gene quantifications, the relative abundance of nitrifiers in seepage was higher in November 2018 than in February and July.

**Figure 6.**
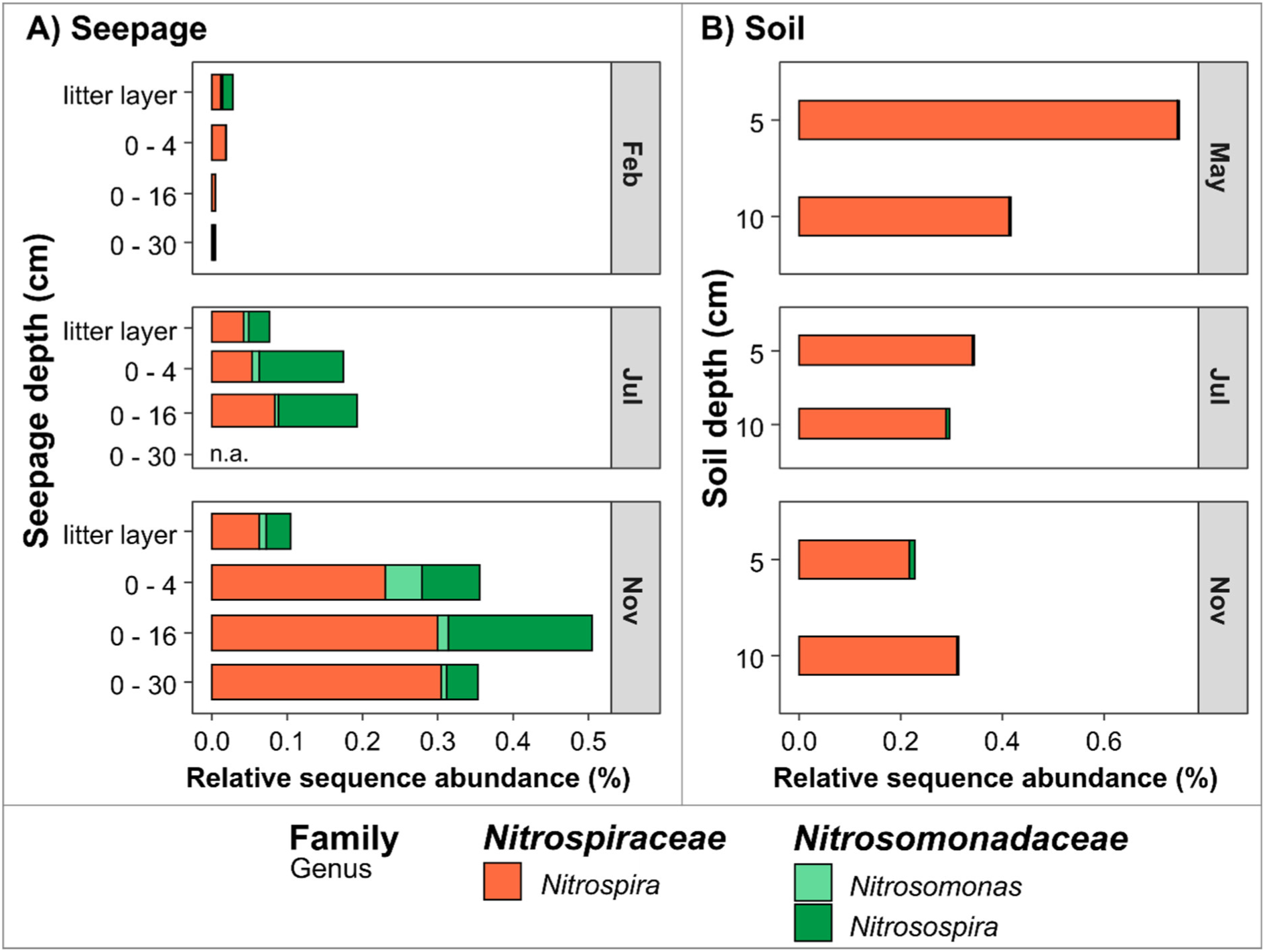
Nitrifier community composition based on 16S rRNA amplicon sequencing of seepage and soil samples. Single bars represent the average relative sequence abundance from all OTUs on genus level assigned to *Nitrospira, Nitrosomonas* and *Nitrosospira* of the corresponding month and depth (n.a. = no sample available).

We analyzed the total bacterial community for which phyla followed a similar depth-dependent trend in their relative abundance in seepage (Figure S4). *Cand*. Patescibacteria showed on average an increasing trend in relative abundance from 6% in seepage below the litter layer to 17% in 30 cm depth, while Bacteroidetes and Verrucomicrobia decreased with increasing seepage depth from 12% and 10% below the litter layer to 2% relative abundance (Figure 7). Moreover, relative abundances varied between time points for every taxon. PCoA analysis indicated a clear differentiation between seepage samples obtained in February 2018 from those obtained in July and November, while no differentiation between sampling time points was observed for the soil samples (Figure S5). In addition, a PERMANOVA approach revealed that sampling months accounted for 17% (*p* < 0.001) and sampling depths accounted for 10% (*p* < 0.05) of the observed variation of bacterial communities in seepage (Table S3).

**Figure 7.**
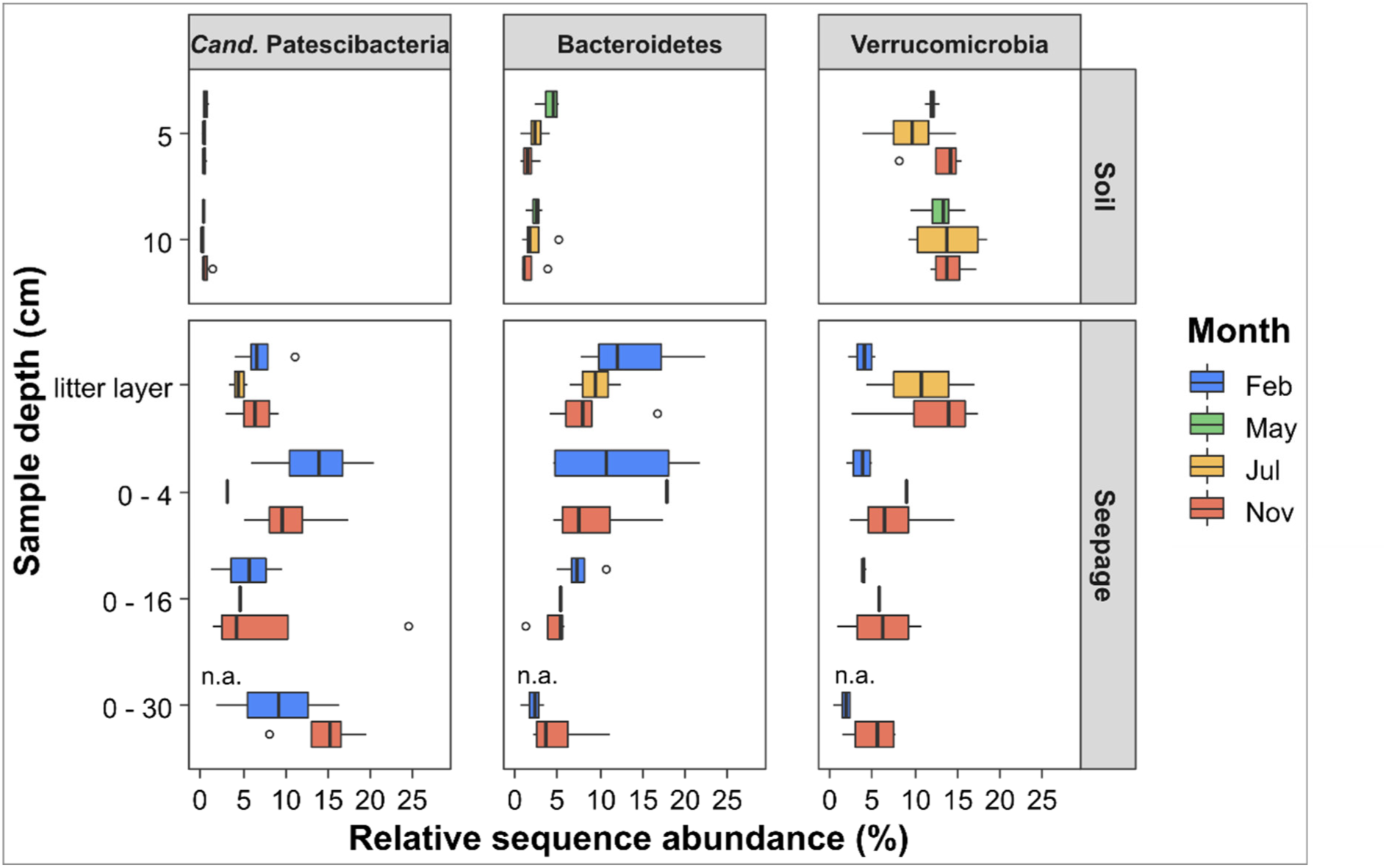
Relative abundances of *Cand*. Patescibacteria, Bacteroidetes and Verrucomicrobia from seepage and soil samples at mixed beech forest plots. Single boxplots represent the average relative sequence abundance from all OTUs of displayed bacterial phyla at particular depths and months (n.a. = no sample available).

## Discussion

Climate models predict an increased frequency of summer drought periods in temperate regions in the future. Before, during and after the severe summer drought in 2018, we aimed to disentangle the relationship between soil nitrification potential, NO_3_ ^-^ leaching and seepage-mediated translocation of soil microorganisms under the impact of summer drought and subsequent autumn precipitation. In contrast to our expectations, the variations of PNR only partially explained the strong NO_3_ ^-^ leaching observed after summer drought and autumn rainfall in the mixed beech forest soils of the Hainich CZE. The highest potential for soil nitrification was measured in soil sampled in May and July during early drought with soil temperatures around 15°C and volumetric water content below 25%. These conditions were closest to optimal conditions for soil nitrification with temperatures between 25 to 30°C and 20% water content (Blume et al., 2010; Carnol and Ineson, 1999; Gilmour, 1984), supporting maximum nitrification activity in July. In addition, fluctuations in NH_4_ ^+^ availability may have contributed to changes in PNR over time. At a concentration of 0.46 µg NH_4_ ^+^ -N g^-1^ soil, nitrification was not limited by NH_4_ ^+^ in July. However, strongly reduced availability of NH_4_ ^+^ in November may have resulted in a lower activity status of the nitrifying populations (Stark and Firestone, 1995), reflected by the lower potential nitrification activity measured in this month.

Overall, the PNR observed in our study ranged from < 0.1 to 4.4 µg N g^-1^ d^-1^, which is comparable to other studies in deciduous (Verchot et al., 2001) and boreal forests (Lu *et al*., 2015). AOB appeared to be more relevant than AOA for soil nitrification activity indicated by higher abundance of bacterial *amoA* genes, and by a positive correlation of bacterial *amoA* genes with PNR and soil NO_3_ ^-^ content. Estimated cell activities of AOB were mostly in the range of previously reported values (Lu et al., 2015; Taylor and Bottomley, 2006), suggesting that activity by AOB alone is sufficient to explain the measured PNR. A predominant role of AOA for nitrification in these soils is unlikely as estimated cell activities exceed commonly reported ranges by one order of magnitude (Lehtovirta-Morley et al., 2011; Lu et al., 2015). This result is in contrast to multiple previous studies which found that AOA are more abundant than AOB in soils and the main driver of nitrification in forest soils (Isobe et al., 2020; Lu et al., 2015) and agricultural soils (Leininger et al., 2006; Sterngren et al., 2015). In particular, an acidic pH below 5 was reported to favor AOA activity rather than AOB (Lu et al., 2015). In the Hainich CZE forest soils, abundances of archaeal *amoA* genes were at least 100 times lower compared to those studies where AOA dominated soil ammonia-oxidizing communities and soil nitrification (Nicol *et al*., 2008; Zhang *et al*., 2012; Lu *et al*., 2015). The observed predominance of AOB over AOA might be explained by the calcareous soils and the parent limestone rock material, as AOB showed a stronger relative contribution to nitrification in calcareous soils (Tao et al., 2017), or after liming (Zhang et al., 2017). Availability of inorganic N and a less acidic pH between 5 to 6 seem to favor AOB which agrees with previous observations that AOB dominate nitrification in soils with high NH_4_ ^+^ supply (Levičnik-Höfferle et al., 2012). Moreover, AOB have higher cell activities due to an overall larger cell volume and faster growth rates (Prosser and Nicol, 2012) and are therefore likely to outcompete AOA under favorable conditions.

Since comammox-related *amoA* gene abundances of *Nitrospira* Clade A and B (Pjevac et al., 2017) remained below the quantification limit for both soil and seepage samples, comammox *Nitrospira* apparently play only a minor role in nitrification at our study site, although a recent study reported a significance role of comammox *Nitrospira* in temperate forest soil nitrification (Osburn and Barrett, 2020).

NO_3_ ^-^ leaching showed more temporal variance than PNR. Highest NO_3_ ^-^ fluxes were observed during summer and autumn 2018, suggesting that NO_3_ ^-^ availability exceeded plant and microbial uptake due to the lower plant N demands after summer (Kaiser et al., 2011). Maxima of NO_3_ ^-^ leaching per sampling event exceeded previous observations from deciduous forests by a factor of two or three (Gundersen et al., 2006; Herrmann et al., 2005). Usually, during autumn and winter months higher amounts of NO_3_ ^-^ are leached out from soils and snow melts further increase NO_3_^-^ leaching in early spring (Herrmann et al., 2005; Jost et al., 2011). NO_3_-leaching can occur as a result of both atmospheric deposition of NO_3_ ^-^ as well as conversion of NH_4_ ^+^ from various sources, including atmospheric deposition, to NO_3_ ^-^ inputs from atmospheric deposition ranging from 1 to 60 kg N ha^-1^ y^-1^ can match NO_3_ ^-^ leaching from European forests soils which ranges between 1 to 40 kg N ha^-1^ y^-1^ (MacDonald et al., 2002).

In the Hainich CZE the average dissolved nitrogen (DN) inputs from throughfall to the soil was reported to be lower than the average DN flux in the soil leachate at 4 cm depth with 12.6 ± 2.1 kg ha^-1^ y^-1^ compared to 22.1 ± 7.3 kg ha^-1^ y^-1^, respectively (Potthast et al., 2017). Soil nitrification is likely favored by NH_4_ ^+^ addition as a portion of DN from throughfall. The ratio of NH_4_ ^+^ to NO_3_ ^-^ is known to decrease from throughfall to soil leachate (Schwarz et al., 2016), suggesting that a fraction of the incoming NH_4_ ^+^ is either immobilized by binding to soil minerals or by uptake into microbial biomass, or is transformed to NO_3_ ^-^ by nitrification. In addition, fluxes of total nitrogen (TN) showed a strong increase from spring to autumn 2018, indicating that drought also enhanced the export of nitrogen from soil (personal communication)

Severe summer droughts, as we observed in the Hainich CZE in 2018, restrains soil N cycling by decreasing plant N uptake and microbial N mineralization, overall reducing microbial activity (Schimel, 2018; Shepherd et al., 2018). In turn, rewetting periods after drought stress fuel soil microbial activity (Hammerl et al., 2019; Placella and Firestone, 2013; Schimel, 2018), cause rapid microbial growth (Göransson et al., 2013; Meisner et al., 2017) and lead to a flush of available N (Homyak et al., 2014). Drought and rewetting can stimulate transcriptional activity of ammonia oxidizers within hours (Placella and Firestone, 2013), suggesting that the low soil NH_4_ ^+^ content we measured in November could be a direct result of increased nitrification stimulated by rainfall in September and October.

Subsequent enhanced NO_3_ ^-^ leaching still continued in November, more than one month after drought conditions had subsided. Leitner et al. (2020) recently reported that summer drought increased NO_3_ ^-^ leaching from forest soils even in the following year after drought. Notably, lower PNR in November compared to May and July indicated that NO_3_ ^-^ leaching in late autumn did not reflect continued intense NO_3_ ^-^ production in the soil but rather mobilization of the NO_3_ ^-^ produced below 4 cm upon rewetting in September and October. In addition, enhanced NO_3_ ^-^ leaching in autumn may further have been supported by the absence of NO_3_ ^-^ sinks, i. e. low N uptake by the forest vegetation at the end of the vegetation period or low NO_3_ ^-^ reduction potential, resulting from a presumably reduced number of anoxic microsites in the well aerated soils after drought. Along with high NO_3_ ^-^ loads, seepage in November after rewetting carried 10 times higher loads of total bacteria as well as AOB and AOA than before or during drought. This observation coincided with a two times higher fraction of POC and PN relative to TOC and TN in seepage during the drought period 2018 and the subsequent autumn months compared to the values before drought and in winter (personal communication), suggesting that drought and subsequent rewetting induced a strong mobilization of particulate organic matter including a substantial fraction of bacteria. This bacterial signal was especially strong below the litter layer and in 4 cm depth but appeared to affect carbon fluxes down to 30 cm depth.

Enhanced microbial growth during rewetting might have contributed to the flush of microbes by infiltrating seepage and to the export of the genetic potential of nitrification towards deeper soil. However, AOB and AOA abundances as well as bacterial abundances in soil showed only minor fluctuations over time, raising the question why increased microbial abundances in seepage were not linked to increased microbial abundances in soil. On the one hand, fast-growing cells might be more susceptible to vertical translocation. Alternatively, a certain fraction of the translocated cells might simply be dead cells which had not survived drought conditions and were now flushed out of the topsoil. Either way, this vertical translocation may have substantial impact on the dynamics and spatial organization of soil microbial communities but also on a potential translocation of microbial carbon to deeper soils and the subsurface.

Moreover, our results provided clear evidence that the susceptibility to vertical translocation is taxon-specific and shows a high degree of differentiation already across the upper 30 cm of the soil. The ratio of soil ammonia oxidizers to their translocated fraction in seepage suggested a stronger vertical translocation of AOA. While AOB were dominating the nitrifier community in 5 to 10 cm of mineral soil, AOA might become more important towards deeper soil, as we calculated that 12% of soil AOA were exported by seepage from upper mineral soil to 30 cm depth. The preferred export of AOA by seepage might be related to a special surface layer (s-layer) covering archaeal cells (Albers and Meyer, 2011). These surface structures consisting of proteins or glycoproteins can increase negative cell surface charge (Sleytr et al., 2014). As soil mineral particles exhibit negative surface charges as well, negatively charged AOA cells could be detached from the soil matrix. Another factor might be the smaller cell size of AOA compared to their bacterial counterparts (Prosser and Nicol, 2012). In contrast, AOB can produce extracellular polymeric substances, consisting of polysaccharides and proteins (Yin et al., 2015), which favor cell aggregation and might prevent detachment from soil particles.

Along with ammonia oxidizers also other bacteria were vertically translocated by seepage. Across all sampling depths, Proteobacteria were by far the most dominant group found in seepage, while Bacteroidetes, Verrucomicrobia and *Cand*. Patescibacteria showed a depth-dependent trend in their translocation patterns. Bacteroidetes and Verrucomicrobia appeared to be most strongly released from the litter layer and the upper 4 cm of soil but retained below 16 cm depth. Attachment in the rhizosphere could be one retention mechanism, as some genera of the phylum Bacteroidetes include plant commensals (Colin et al., 2017). The small-sized cells of *Cand*. Patescibacteria, with a maximum of mobilization below the litter layer to 16 and 30 cm depth, were not retained and can be exported to subsurface environments like groundwater (Herrmann et al., 2019). This preferential transport of microbes also showed variation between different months. Lower seepage volumes and drier soils in summer, especially under drought conditions, appear to prevent mobilization of *Cand*. Patescibacteria, while more cells were translocated during autumn and winter. Rewetting also seemed to stimulate the mobilization of Verrucomicrobia via seepage in November, while Bacteroidetes were not affected. These observed taxon-specific translocation patterns of selected topsoil derived microbial groups imply a selective transfer of genetic functions for biogeochemical processes and nutrient cycling from near surface soils to subsoils.

## Conclusions

Our results clearly demonstrated that summer drought followed by autumn precipitation led to strong nitrate leaching from soils of a mixed beech forest. Temporal dynamics of leaching were only partially coupled to temporal dynamics of soil potential nitrification activity, suggesting that not only enhanced nitrate production during summer but also missing nitrate sinks in dry soils contributed to the strong autumn nitrate leaching. Increased nitrate fluxes after drought and rewetting coincided with a 10 times higher translocation of ammonia oxidizing bacteria and archaea as well as total bacteria with seepage across the upper 30 cm of the soil, which reflected intense microbial growth after rewetting but probably also export of dead cells in response to the extreme conditions. Given the increase of drought periods in temperate regions based on future climate models, the observed drought and rewetting effects on nitrate leaching and bacterial translocation may occur more frequently. Moreover, microbial translocation patterns pointed to taxon-specific differences in the susceptibility to mobilization, suggesting that only selected groups are likely to affect subsoil or groundwater microbial communities and their functional potential.

## Acknowledgements

We greatly thank Marie-Cecile Gruselle, Stefan Krümmling, Stefan Riedel, Valentin Kurbel, Lena Carstens and Beatrix Heinze for field and laboratory assistance. We further thank Maria Fabisch for scientific coordination within the CRC AquaDiva framework.

This study was part of the Collaborative Research Centre 1076 AquaDiva (CRC AquaDiva) of the Friedrich Schiller University Jena, funded by the Deutsche Forschungsgemeinschaft. Climate chambers to conduct experiments under controlled temperature conditions and the infrastructure for MiSeq Illumina sequencing were financially supported by the Thüringer Ministerium für Wirtschaft, Wissenschaft und Digitale Gesellschaft (TMWWDG; project B 715-09075 and project 2016 FGI 0024 “BIODIV”).

## Author contributions

MK carried out potential nitrification rate measurements and most of the chemical analyses and molecular work. MH and MK planned the project. BM and KP designed and installed the field infrastructure. KP and AT provided results of chemical analyses of seepage. FFKD contributed to field work, chemical analysis, and molecular work. MK, MH and KK wrote the manuscript with contributions from all other authors.

## References

Abu-Ashour, J., Joy, D.M., Lee, H., Whiteley, H.R., Zelin, S., 1994. Transport of microorganisms through soil. Water, Air, & Soil Pollution 75, 141–158.

Albers, S.V., Meyer, B.H., 2011. The archaeal cell envelope. Nature Reviews Microbiology.

Blume, H.-P., Brümmer, G.W., Horn, R., Kandeler, E., Kögel-Knabner, I., Kretzschmar, R., Stahr, K., Wilke, B.-M., Thiele-Bruhn, S., Welp, G., 2010. Lehrbuch der Bodenkunde, Springer.

Borken, W., Matzner, E., 2004. Nitrate leaching in forest soils: an analysis of long-term monitoring sites in Germany. Journal of Plant Nutrition and Soil Science 167, 277–283.

Carnol, M., Ineson, P., 1999. Environmental factors controlling NO3-leaching, N2O emissions and numbers of NH4+ oxidisers in a coniferous forest soil. Soil Biology and Biochemistry 31, 979–990.

Colin, Y., Nicolitch, O., Van Nostrand, J.D., Zhou, J.Z., Turpault, M.P., Uroz, S., 2017. Taxonomic and functional shifts in the beech rhizosphere microbiome across a natural soil toposequence. Scientific Reports 7, 1–17.

Corre, M.D., Beese, F.O., Brumme, R., 2003. Soil nitrogen cycle in high nitrogen deposition forest: changes under nitrogen saturation and liming. Ecological Applications 13, 287–298.

Daims, H., Brühl, A., Amann, R., Schleifer, K.H., Wagner, M., 1999. The domain-specific probe EUB338 is insufficient for the detection of all bacteria: development and evaluation of a more comprehensive probe set. Systematic and Applied Microbiology 22, 434–444.

Dibbern, D., Schmalwasser, A., Lueders, T., Totsche, K.U., 2014. Selective transport of plant root-associated bacterial populations in agricultural soils upon snowmelt. Soil Biology and Biochemistry 69, 187–196.

Edgar, R.C., Haas, B.J., Clemente, J.C., Quince, C., Knight, R., 2011. UCHIME improves sensitivity and speed of chimera detection. Bioinformatics 27, 2194–2200.

Francis, C.A., Roberts, K.J., Beman, J.M., Santoro, A.E., Oakley, B.B., 2005. Ubiquity and diversity of ammonia-oxidizing archaea in water columns and sediments of the ocean. Proceedings of the National Academy of Sciences 102, 14683–14688.

Gilmour, J.T., 1984. The effects of soil properties on nitrification and nitrification inhibition 1262–1267.

Göransson, H., Godbold, D.L., Jones, D.L., Rousk, J., 2013. Bacterial growth and respiration responses upon rewetting dry forest soils: impact of drought-legacy. Soil Biology and Biochemistry 57, 477–486.

Gordon, H., Haygarth, P.M., Bardgett, R.D., 2008. Drying and rewetting effects on soil microbial community composition and nutrient leaching. Soil Biology and Biochemistry 40, 302–311.

Gruber, N., Galloway, J.N., 2008. An Earth-system perspective of the global nitrogen cycle. Nature.

Gundersen, P., Schmidt, I.K., Raulund-Rasmussen, K., 2006. Leaching of nitrate from temperate forests - effects of air pollution and forest management. Environmental Reviews 14, 1–57.

Hallam, S.J., Konstantinidis, K.T., Putnam, N., Schleper, C., Watanabe, Y.I., Sugahara, J., Preston, C., De La Torre, J., Richardson, P.M., DeLong, E.F., 2006. Genomic analysis of the uncultivated marine crenarchaeote Cenarchaeum symbiosum. Proceedings of the National Academy of Sciences of the United States of America 103, 18296–18301.

Hammerl, V., Kastl, E.M., Schloter, M., Kublik, S., Schmidt, H., Welzl, G., Jentsch, A., Beierkuhnlein, C., Gschwendtner, S., 2019. Influence of rewetting on microbial communities involved in nitrification and denitrification in a grassland soil after a prolonged drought period. Scientific Reports 9, 2280.

Hart, C.S., Stark, M.J., Davidson, A.E., Firestone, K.M., 1994. Nitrogen mineralization, immobilization, and nitrification.

Herlemann, D.P.R., Labrenz, M., Jürgens, K., Bertilsson, S., Waniek, J.J., Andersson, A.F., 2011. Transitions in bacterial communities along the 2000 km salinity gradient of the Baltic Sea. The ISME Journal 5, 1571–1579.

Herrmann, M., Hädrich, A., Küsel, K., 2012. Predominance of thaumarchaeal ammonia oxidizer abundance and transcriptional activity in an acidic fen. Environmental Microbiology 14, 3013–3025.

Herrmann, M., Pust, J., Pott, R., 2005. Leaching of nitrate and ammonium in heathland and forest ecosystems in Northwest Germany under the influence of enhanced nitrogen deposition. Plant and Soil 273, 129–137.

Herrmann, M., Wegner, C.-E., Taubert, M., Geesink, P., Lehmann, K., Yan, L., Lehmann, R., Totsche, K.U., Küsel, K., 2019. Predominance of Cand. Patescibacteria in groundwater is caused by their preferential mobilization from soils and flourishing under oligotrophic conditions. Frontiers in Microbiology 10, 1407.

Homyak, P.M., Sickman, J.O., Miller, A.E., Melack, J.M., Meixner, T., Schimel, J.P., 2014. Assessing nitrogen-saturation in a seasonally dry chaparral watershed: limitations of traditional indicators of N-saturation. Ecosystems 17, 1286–1305.

Hu, H.W., He, J.Z., 2017. Comammox—a newly discovered nitrification process in the terrestrial nitrogen cycle. Journal of Soils and Sediments.

Isobe, K., Ise, Y., Kato, H., Oda, T., Vincenot, C.E., Koba, K., Tateno, R., Senoo, K., Ohte, N., 2020. Consequences of microbial diversity in forest nitrogen cycling: diverse ammonifiers and specialized ammonia oxidizers. The ISME Journal 14, 12–25.

Jost, G., Dirnböck, T., Grabner, M.T., Mirtl, M., 2011. Nitrogen leaching of two forest ecosystems in a karst watershed. Water, Air, and Soil Pollution 218, 633–649.

Kaiser, C., Fuchslueger, L., Koranda, M., Gorfer, M., Stange, C.F., Kitzler, B., Rasche, F., Strauss, J., Sessitsch, A., Zechmeister-Boltenstern, S., Richter, A., 2011. Plants control the seasonal dynamics of microbial N cycling in a beech forest soil by belowground C allocation. Ecology 92, 1036–1051.

Könneke, M., Bernhard, A.E., De La Torre, J.R., Walker, C.B., Waterbury, J.B., Stahl, D.A., 2005. Isolation of an autotrophic ammonia-oxidizing marine archaeon. Nature 437, 543–546.

Kowalchuk, G.A., Stephen, J.R., 2001. Ammonia-oxidizing bacteria: a model for molecular microbial ecology. Annual Review of Microbiology 55, 485–529.

Kozich, J.J., Westcott, S.L., Baxter, N.T., Highlander, S.K., Schloss, P.D., 2013. Development of a dual-index sequencing strategy and curation pipeline for analyzing amplicon sequence data on the miseq illumina sequencing platform. Applied and Environmental Microbiology 79, 5112–5120.

Küsel, K., Totsche, K.U., Trumbore, S.E., Lehmann, R., Steinhäuser, C., Herrmann, M., 2016. How deep can surface signals be traced in the Critical Zone? Merging biodiversity with biogeochemistry research in a central German Muschelkalk landscape. Frontiers in Earth Science 4, 32.

Lehmann, K., Schaefer, S., Babin, D., Köhne, J.M., Schlüter, S., Smalla, K., Vogel, H.J., Totsche, K.U., 2018. Selective transport and retention of organic matter and bacteria shapes initial pedogenesis in artificial soil - a two-layer column study. Geoderma 325, 37–48.

Lehmann, R., Totsche, K.U., 2020. Multi-directional flow dynamics shape groundwater quality in sloping bedrock strata. Journal of Hydrology 580, 124291.

Lehtovirta-Morley, L.E., 2018. Ammonia oxidation: ecology, physiology, biochemistry and why they must all come together. FEMS Microbiology Letters 365.

Lehtovirta-Morley, L.E., Stoecker, K., Vilcinskas, A., Prosser, J.I., Nicol, G.W., 2011. Cultivation of an obligate acidophilic ammonia oxidizer from a nitrifying acid soil. Proceedings of the National Academy of Sciences 108, 15892–15897.

Leininger, S., Urich, T., Schloter, M., Schwark, L., Qi, J., Nicol, G.W., Prosser, J.I., Schuster, S.C., Schleper, C., 2006. Archaea predominate among ammonia-oxidizing prokaryotes in soils. Nature 442, 806–809.

Leitner, S., Dirnböck, T., Kobler, J., Zechmeister-Boltenstern, S., 2020. Legacy effects of drought on nitrate leaching in a temperate mixed forest on karst. Journal of Environmental Management 262.

Levičnik-Höfferle, Š., Nicol, G.W., Ausec, L., Mandić-Mulec, I., Prosser, J.I., 2012. Stimulation of thaumarchaeal ammonia oxidation by ammonia derived from organic nitrogen but not added inorganic nitrogen. FEMS Microbiology Ecology 80, 114–123.

Loy, A., Lehner, A., Lee, N., Adamczyk, J., Meier, H., Ernst, J., Schleifer, K.-H., Wagner, M., 2002. Oligonucleotide microarray for 16S rRNA gene-based detection of all recognized lineages of sulfate-reducing prokaryotes in the environment. Applied and Environmental Microbiology 68, 5064–5081.

Lu, X., Bottomley, P.J., Myrold, D.D., 2015. Contributions of ammonia-oxidizing archaea and bacteria to nitrification in Oregon forest soils. Soil Biology and Biochemistry 85, 54–62.

Luef, B., Frischkorn, K.R., Wrighton, K.C., Holman, H.Y.N., Birarda, G., Thomas, B.C., Singh, A., Williams, K.H., Siegerist, C.E., Tringe, S.G., Downing, K.H., Comolli, L.R., Banfield, J.F., 2015. Diverse uncultivated ultra-small bacterial cells in groundwater. Nature Communications 6, 1–8.

MacDonald, J.A., Dise, N.B., Matzner, E., Armbruster, M., Gundersen, P., Forsius, M., 2002. Nitrogen input together with ecosystem nitrogen enrichment predict nitrate leaching from European forests. Global Change Biology 8, 1028–1033.

McMurdie, P.J., Holmes, S., 2013. Phyloseq: an R package for reproducible interactive analysis and graphics of microbiome census data. PLoS ONE 8, e61217.

Meisner, A., Leizeaga, A., Rousk, J., Bååth, E., 2017. Partial drying accelerates bacterial growth recovery to rewetting. Soil Biology and Biochemistry 112, 269–276.

Neff, J.C., III, F.S.C., Vitousek, P.M., 2003. Breaks in the cycle: dissolved organic nitrogen in terrestrial ecosystems. Frontiers in Ecology and the Environment 1, 205.

Nicol, G.W., Leininger, S., Schleper, C., Prosser, J.I., 2008. The influence of soil pH on the diversity, abundance and transcriptional activity of ammonia oxidizing archaea and bacteria. Environmental Microbiology 10, 2966–2978.

Norton, J.M., Alzerreca, J.J., Suwa, Y., Klotz, M.G., 2002. Diversity of ammonia monooxygenase operon in autotrophic ammonia-oxidizing bacteria. Archives of Microbiology 177, 139–149.

Oksanen, J., Kindt, R., Legendre, P., O’Hara, B., Simpson, G.L., Solymos, P.M., Stevens, M.H.H., & Wagner, H., 2008. The vegan package. Community Ecology Package 190.

Osburn, E.D., Barrett, J.E., 2020. Abundance and functional importance of complete ammonia-oxidizing bacteria (comammox) versus canonical nitrifiers in temperate forest soils. Soil Biology and Biochemistry 145, 2015–2018.

Pjevac, P., Schauberger, C., Poghosyan, L., Herbold, C.W., van Kessel, M.A.H.J., Daebeler, A., Steinberger, M., Jetten, M.S.M., Lücker, S., Wagner, M., Daims, H., 2017. AmoA-targeted polymerase chain reaction primers for the specific detection and quantification of comammox Nitrospira in the environment. Frontiers in Microbiology 8, 1508.

Placella, S.A., Firestone, M.K., 2013. Transcriptional response of nitrifying communities to wetting of dry soil. Applied and Environmental Microbiology 79, 3294–3302.

Potthast, K., Meyer, S., Crecelius, A.C., Schubert, U.S., Tischer, A., Michalzik, B., 2017. Land-use and fire drive temporal patterns of soil solution chemistry and nutrient fluxes. Science of The Total Environment 605–606, 514–526.

Prosser, J.I., 2014. Soil nitrifiers and nitrification. Nitrification 347–383.

Prosser, J.I., Nicol, G.W., 2012. Archaeal and bacterial ammonia-oxidisers in soil: the quest for niche specialisation and differentiation. Trends in Microbiology.

Quast, C., Pruesse, E., Yilmaz, P., Gerken, J., Schweer, T., Yarza, P., Peplies, J., Glöckner, F.O., 2013. The SILVA ribosomal RNA gene database project: improved data processing and web-based tools. Nucleic Acids Research 41, 590–596.

Reay, D.S., Dentener, F., Smith, P., Grace, J., Feely, R.A., 2008. Global nitrogen deposition and carbon sinks. Nature Geoscience 1, 430–437.

Rotthauwe, J.H., Witzel, K.P., Liesack, W., 1997. The ammonia monooxygenase structural gene amoA as a functional marker: molecular fine-scale analysis of natural ammonia-oxidizing populations. Applied and Environmental Microbiology 63, 4704–12.

Schaeffer, S.M., Homyak, P.M., Boot, C.M., Roux-Michollet, D., Schimel, J.P., 2017. Soil carbon and nitrogen dynamics throughout the summer drought in a California annual grassland. Soil Biology and Biochemistry 115, 54–62.

Scheiner, D., 1974. A modified version of the sodium salicylate method for analysis of wastewater nitrates. Water Research 8, 835–840.

Schimel, J., Balser, T.C., Wallenstein, M., 2007. Microbial stress-response physiology and its implications for ecosystem function. Ecology 88, 1386–1394.

Schimel, J.P., 2018. Life in dry soils: effects of drought on soil microbial communities and processes. Annual Review of Ecology, Evolution, and Systematics 49, 409–432.

Schlesinger, W.H., 2009. On the fate of anthropogenic nitrogen. Proceedings of the National Academy of Sciences of the United States of America 106, 203–208.

Schlesinger, W.H., Dietze, M.C., Jackson, R.B., Phillips, R.P., Rhoades, C.C., Rustad, L.E., Vose, J.M., 2016. Forest biogeochemistry in response to drought. Global Change Biology 22, 2318–2328.

Schloss, P.D., Westcott, S.L., Ryabin, T., Hall, J.R., Hartmann, M., Hollister, E.B., Lesniewski, R.A., Oakley, B.B., Parks, D.H., Robinson, C.J., Sahl, J.W., Stres, B., Thallinger, G.G., Van Horn, D.J., Weber, C.F., 2009. Introducing mothur: open-source, platform-independent, community-supported software for describing and comparing microbial communities. Applied and Environmental Microbiology 75, 7537–7541.

Schwarz, M.T., Bischoff, S., Blaser, S., Boch, S., Grassein, F., Klarner, B., Schmitt, B., Solly, E.F., Ammer, C., Michalzik, B., Schall, P., Scheu, S., Schöning, I., Schrumpf, M., Schulze, E.D., Siemens, J., Wilcke, W., 2016. Drivers of nitrogen leaching from organic layers in central European beech forests. Plant and Soil 403, 343–360.

Seelmeyer, G., 1954. Deutsche Einheitsverfahren zur Wasseruntersuchung. Herausgegeben im Auftrage der Fachgruppe Wasserchemie inder Gesellschaft Deutscher Chemiker E. V. Bearbeitet von L. W. Haase, H. Stooff, G. Gad, W. Wesly u.a. Verlag Chemie, Weinheim/Bergstraße 1954, 180. Materials and Corrosion 5, 357–357.

Shepherd, M., Lucci, G., Vogeler, I., Balvert, S., 2018. The effect of drought and nitrogen fertiliser addition on nitrate leaching risk from a pasture soil; an assessment from a field experiment and modelling. Journal of the Science of Food and Agriculture 98, 3795–3805.

Sleytr, U.B., Schuster, B., Egelseer, E.-M., Pum, D., 2014. S-layers: principles and applications. FEMS Microbiology Reviews 38, 823–864.

Stark, J.M., Firestone, M.K., 1995. Mechanisms for soil moisture effects on activity of nitrifying bacteria. Applied and Environmental Microbiology 61, 218–221.

Sterngren, A.E., Hallin, S., Bengtson, P., 2015. Archaeal ammonia oxidizers dominate in numbers, but bacteria drive gross nitrification in n-amended grassland soil. Frontiers in Microbiology 6, 1350.

Stoddard, S.F., Smith, B.J., Hein, R., Roller, B.R.K., Schmidt, T.M., 2015. rrnDB: Improved tools for interpreting rRNA gene abundance in bacteria and archaea and a new foundation for future development. Nucleic Acids Research 43, D593–D598.

Tao, R., Wakelin, S.A., Liang, Y., Chu, G., 2017. Response of ammonia-oxidizing archaea and bacteria in calcareous soil to mineral and organic fertilizer application and their relative contribution to nitrification. Soil Biology and Biochemistry 114, 20–30.

Taylor, A.E., Bottomley, P.J., 2006. Nitrite production by Nitrosomonas europaea and Nitrosospira sp. AV in soils at different solution concentrations of ammonium. Soil Biology and Biochemistry 38, 828–836.

Van Kessel, M.A.H.J., Speth, D.R., Albertsen, M., Nielsen, P.H., Op Den Camp, H.J.M., Kartal, B., Jetten, M.S.M., Lücker, S., 2015. Complete nitrification by a single microorganism. Nature 528, 555–559.

Verchot, L. V., Holmes, Z., Mulon, L., Groffman, P.M., Lovett, G.M., 2001. Gross vs net rates of N mineralization and nitrification as indicators of functional differences between forest types. Soil Biology and Biochemistry 33, 1889–1901.

Vrede, K., Heldal, M., Norland, S., Bratbak, G., 2002. Elemental composition (C, N, P) and cell volume of exponentially growing and nutrient-limited bacterioplankton. Applied and Environmental Microbiology 68, 2965–2971.

Yao, H., Gao, Y., Nicol, G.W., Campbell, C.D., Prosser, J.I., Zhang, L., Han, W., Singh, B.K., 2011. Links between ammonia oxidizer community structure, abundance, and nitrification potential in acidic soils. Applied and Environmental Microbiology 77, 4618–4625.

Yin, C., Meng, F., Chen, G.H., 2015. Spectroscopic characterization of extracellular polymeric substances from a mixed culture dominated by ammonia-oxidizing bacteria. Water Research 68, 740–749.

Zhang, L., Lehmann, K., Totsche, K.U., Lueders, T., 2018. Selective successional transport of bacterial populations from rooted agricultural topsoil to deeper layers upon extreme precipitation events. Soil Biology and Biochemistry 124, 168–178.

Zhang, L.M., Hu, H.W., Shen, J.P., He, J.Z., 2012. Ammonia-oxidizing archaea have more important role than ammonia-oxidizing bacteria in ammonia oxidation of strongly acidic soils. ISME Journal 6, 1032–1045.

Zhang, M.-M., Alves, R.J.E., Zhang, D.-D., Han, L.-L., He, J.-Z., Zhang, L.-M., 2017. Time-dependent shifts in populations and activity of bacterial and archaeal ammonia oxidizers in response to liming in acidic soils. Soil Biology and Biochemistry 112, 77–89.

Zhu, X., Zhang, W., Chen, H., Mo, J., 2015. Impacts of nitrogen deposition on soil nitrogen cycle in forest ecosystems: A review. Acta Ecologica Sinica 35, 35–43.

