## Supplemental figures and tables for "Drought and rewetting events enhance nitrate leaching and seepage-mediated translocation of microbes from beech forest soils"

### Supplementary material

Markus Krüger^1^, Karin Potthast^2^, Beate Michalzik^2,3^, Alexander Tischer^2^, Kirsten Küsel^1,3^, Florian F. K. Deckner^1^, Martina Herrmann^1,3,*^

#### Figure S1

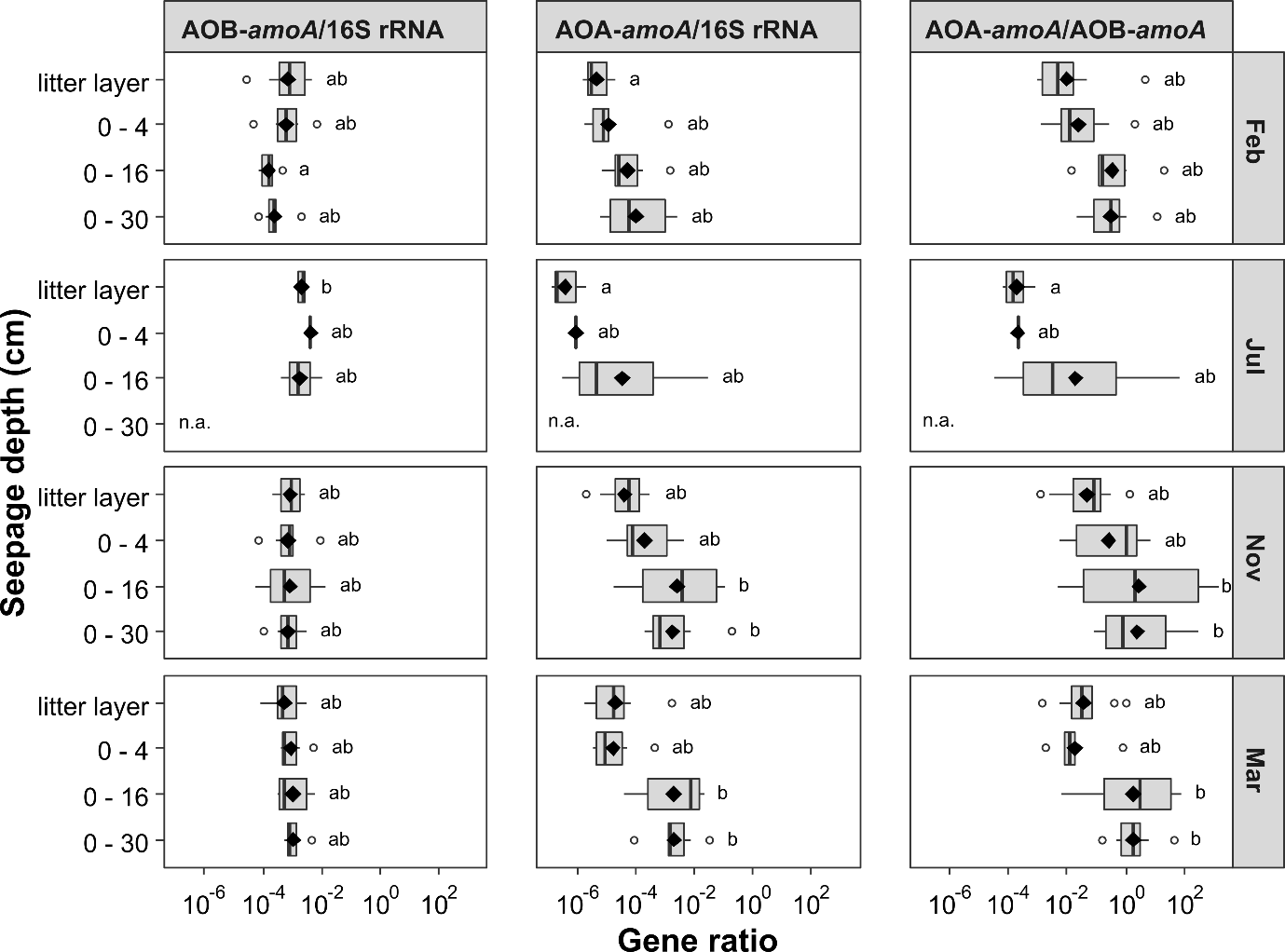

Figure S1. Calculated gene ratios of bacterial *amoA*/16S rRNA, archaeal *amoA*/16S rRNA and archaeal *amoA*/bacterial *amoA* gene abundance in different seepage depths. Individual boxplots summarize the gene ratios at particular depths and black diamonds represent mean of respective data (n.a. = no sample available). The lowercase letter code indicates significant differences based on Dunn’s multiple comparison test (*p* < 0.05).

#### Figure S2

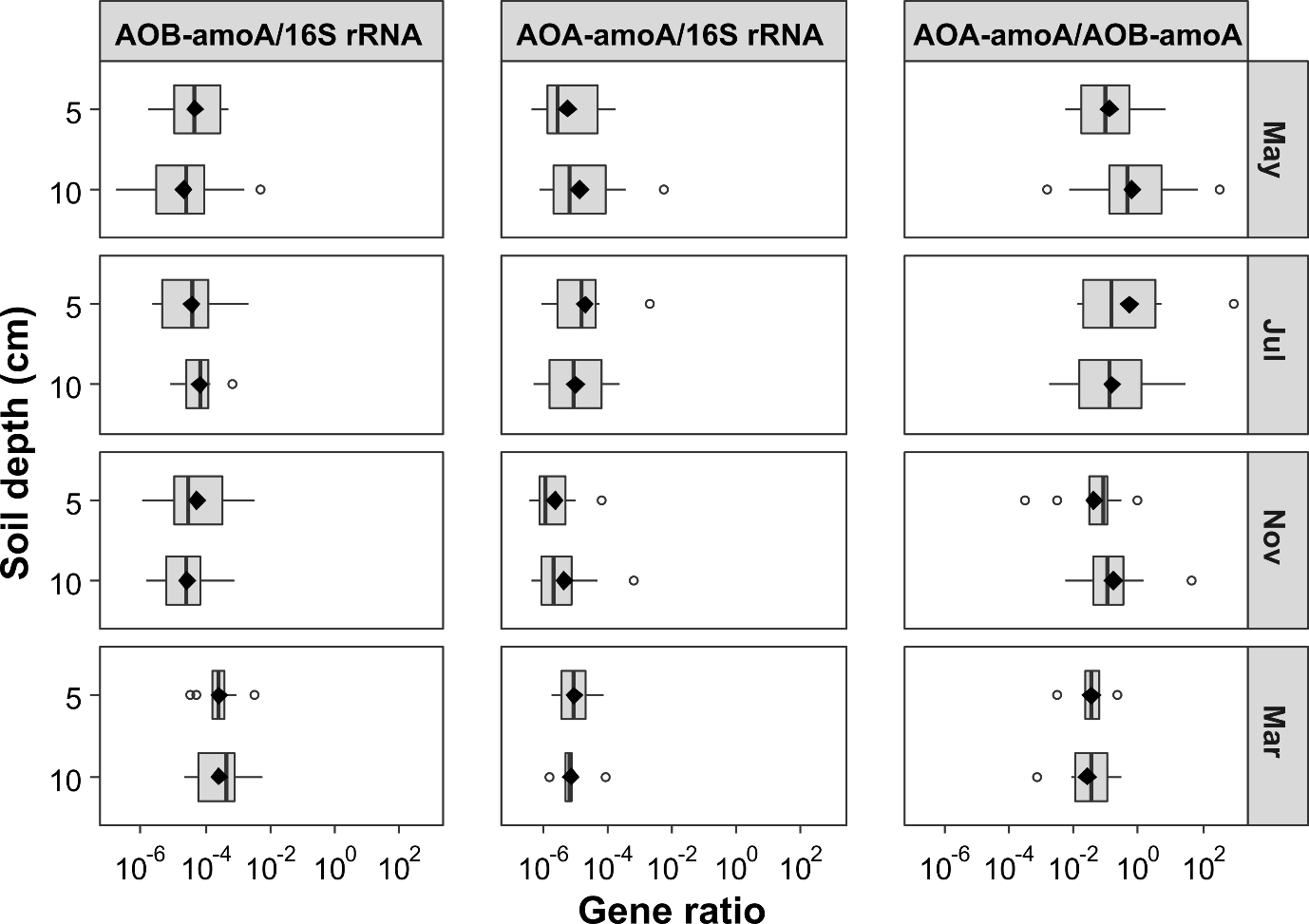

Figure S2. Calculated gene ratios of bacterial *amoA* and archaeal *amoA* to bacterial 16S rRNA gene abundance, and ratios of archaeal *amoA* to bacterial *amoA* gene abundance in different soil depths.

#### Figure S3

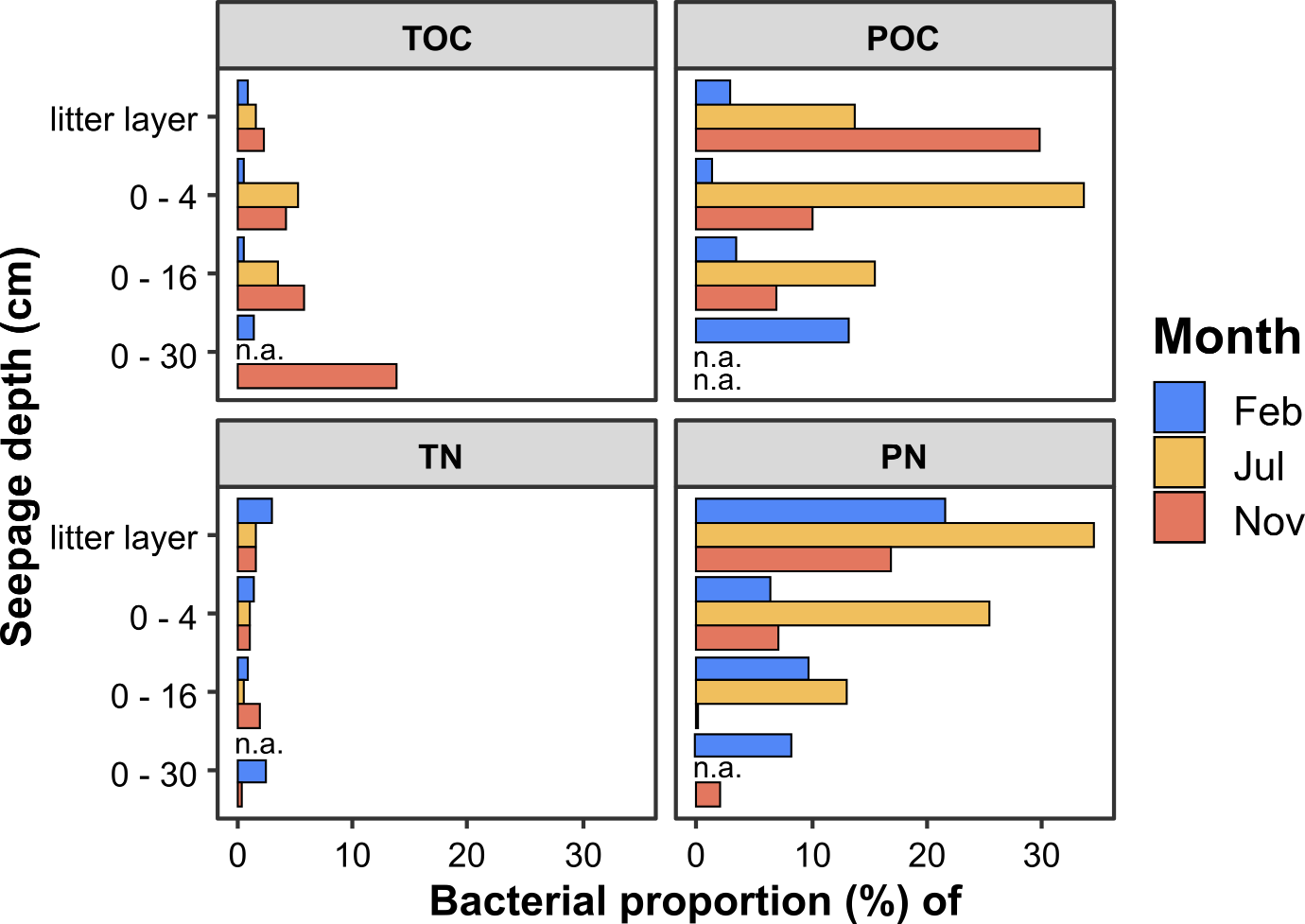

Figure S3. Estimated proportions of bacterial cells to total organic carbon (TOC), particulate organic carbon (POC), total nitrogen (TN) and particulate nitrogen (PN) of seepage samples from February, July and November 2018 from four lysimeter depths. Each bar represents the mean proportion at a particular depth and month (n.a. = no measurement available).

#### Figure S4

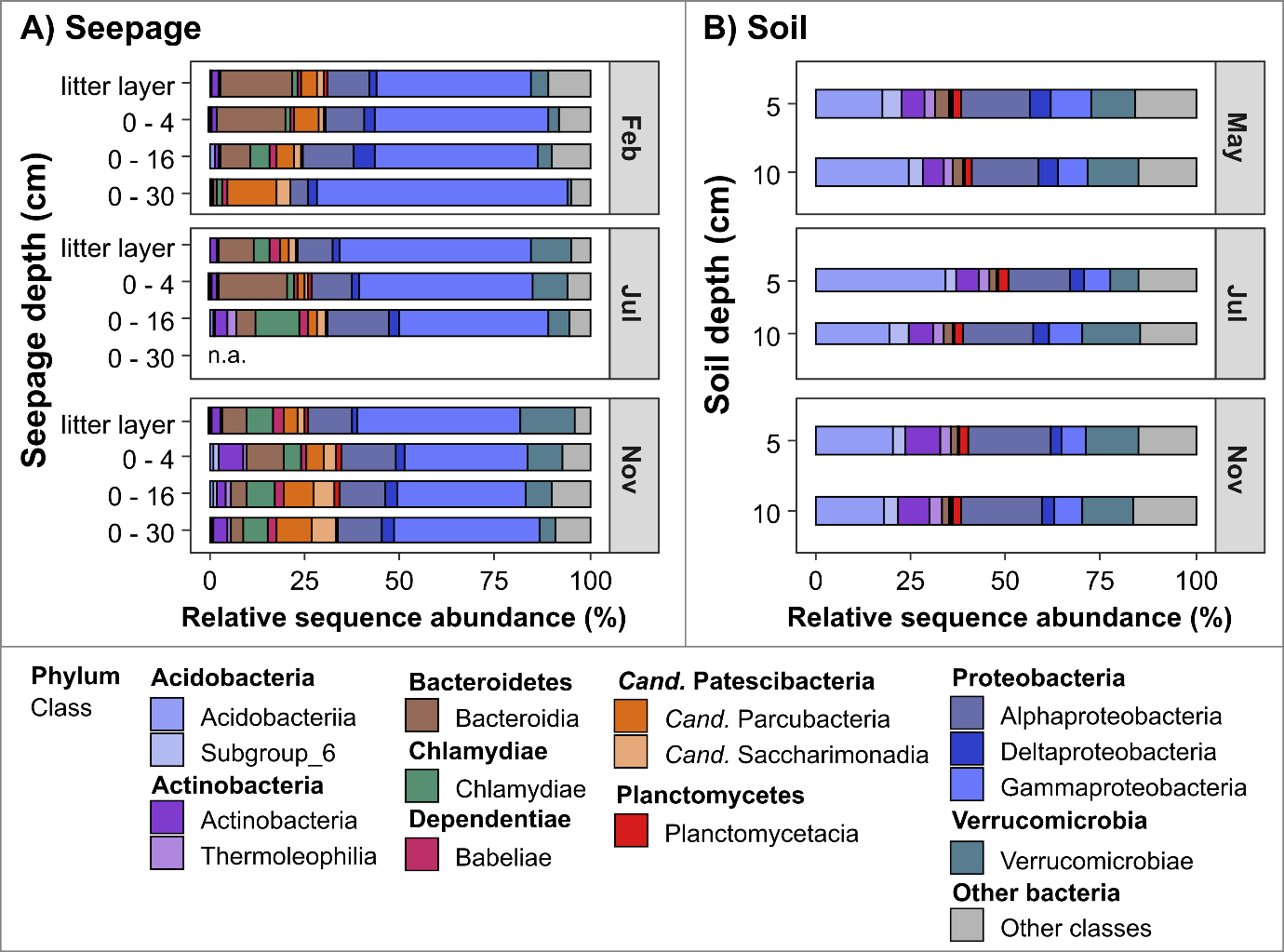

Figure S4. Bacterial community composition based on 16S rRNA amplicon sequencing of A) seepage and B) soil samples. Single bars represent the average relative sequence abundance from the most abundant bacteria (above 1% per sample) on class level of the corresponding months and depths (n.a. = no sample available).

#### Figure S5

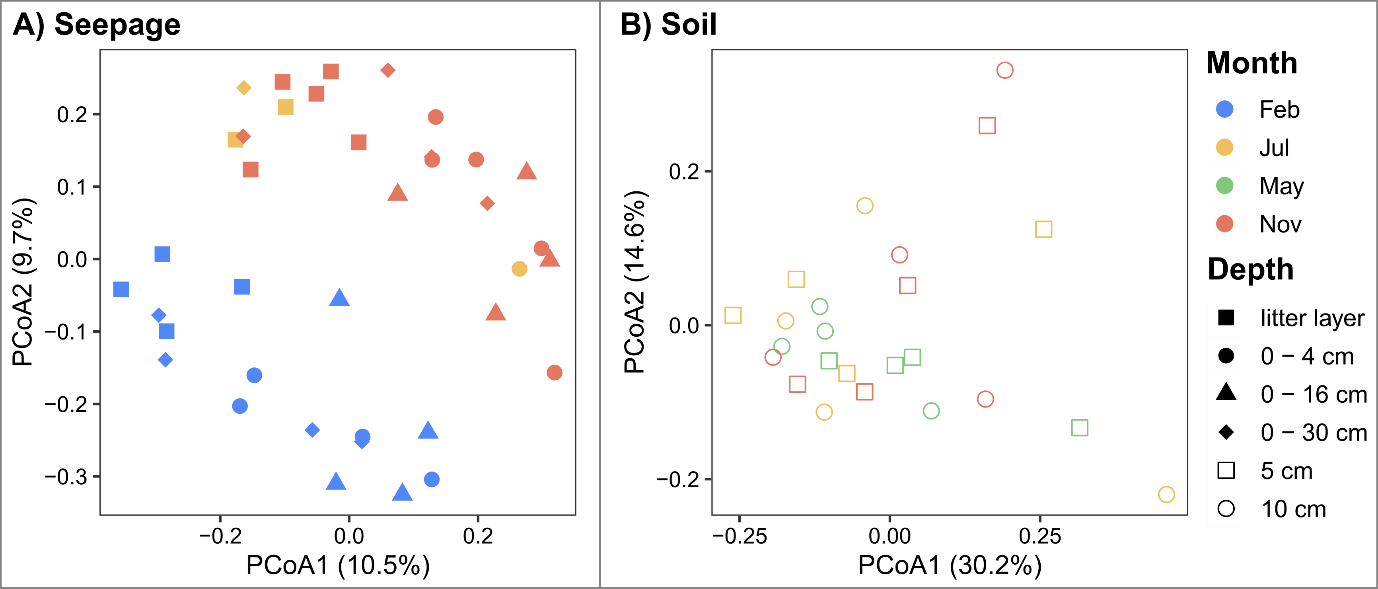

Figure S5. Principal coordinate analysis (PCoA) of OTUs derived from 16S rRNA amplicon sequencing displays clustering of A) seepage of four lysimeter depths and B) soil samples of two depths from samples collected in February, May, July, and November 2018.

#### Table S1

Table S1. Overview of soil temperature and soil moisture sensors placed at 4 cm, 16 cm, and 30 cm depth in mixed beech forest soils. Shown are average values and standard deviation (in parentheses) of a 14 days’ time span before sampling in respective months.

|  |  | **Temperature (°C)** | | |  | **Moisture (%)** | | | | |
| --- | --- | --- | --- | --- | --- | --- | --- | --- | --- | --- |
| **Month** |  | **4 cm** |  | **16 cm** |  | **4 cm** |  | **16 cm** |  | **30 cm** |
| **February** |  | 1.7 (1.1) |  | 2.3 (1.1) |  | 50.6 (3.3) |  | 39.5 (1.2) |  | 43.9 (1.6) |
| **May** |  | 10.7 (1.2) |  | 10.3 (0.8) |  | 37.0 (3.0) |  | 38.0 (2.5) |  | 43.4 (1.6) |
| **July** |  | 14.6 (1.0) |  | 14.2 (0.6) |  | 14.5 (2.2) |  | 20.9 (1.4) |  | 35.4 (2.1) |
| **November** |  | 8.8 (1.1) |  | 9.3 (1.0) |  | 13.7 (0.9) |  | 20.5 (1.4) |  | 36.2 (1.4) |
| **March** |  | 4.6 (1.0) |  | 4.7 (0.8) |  | 32.0 (3.4) |  | 35.8 (1.7) |  | 43.1 (1.2) |

#### Table S2

Table S2. Pairwise correlation analysis based on spearman’s rank correlation coefficient between soil properties of PNR (potential nitrification rate), NO_3_^-^ content, NH_4_^+^ content, Moist (soil moisture), temperature, pH and microbial gene abundance of AOB (bacterial *amoA*) and AOA (archaeal *amoA*). Upper section displays spearman’s rank coefficient and lower section shows significance of observed correlation (ns = not significant, * *p* < 0.05, *** *p* < 0.001, **** *p* < 0.0001).

|  | **PNR** | **NO_3_^-^** | **NH_4_^+^** | **Moist** | **Temp** | **pH_(H2O)_** | **AOB** | **AOA** |
| --- | --- | --- | --- | --- | --- | --- | --- | --- |
| **PNR** |  | 0.28 | 0.34 | 0.18 | 0.01 | 0.27 | 0.56 | 0.31 |
| **NO_3_^-^** | *** |  | -0.31 | -0.31 | -0.10 | -0.06 | 0.34 | 0.14 |
| **NH_4_^+^** | **** | **** |  | 0.36 | 0.13 | 0.43 | 0.20 | 0.19 |
| **Moist** | ns | ** | ** |  | -0.63 | 0.27 | -0.08 | 0.15 |
| **Temp** | ns | ns | ns | **** |  | -0.23 | -0.05 | -0.05 |
| **pH_(H2O)_** | * | ns | **** | ns | ns |  | 0.62 | 0.21 |
| **AOB** | **** | ** | ns | ns | ns | *** |  | 0.28 |
| **AOA** | ** | ns | ns | ns | ns | ns | ** |  |

#### Table S3

Table S3. Permutational multivariant analysis of variance (PERMANOVA) of factors accounting for OTU variances across sampling months and depths from seepage bacterial communities. Default 999 permutations were assessed based on Bray-Curtis distance using R package vegan.

|  | **Df** | **SumsOfSqs** | **MeanSqs** | **F,Model** | **R2** | **Pr(>F)** |
| --- | --- | --- | --- | --- | --- | --- |
| **Month** | 2 | 2.54 | 1.27 | 3.73 | 0.17 | 0.001 |
| **Depth** | 3 | 1.43 | 0.48 | 1.40 | 0.10 | 0.012 |
| **Residuals** | 32 | 10.87 | 0.34 | - | 0.73 | - |
| **Total** | 37 | 14.84 | - | - | 1.00 | - |
